# Structure of Gcn1 bound to stalled and colliding 80S ribosomes

**DOI:** 10.1101/2020.10.31.363135

**Authors:** Agnieszka A. Pochopien, Bertrand Beckert, Sergo Kasvandik, Otto Berninghausen, Roland Beckmann, Tanel Tenson, Daniel N. Wilson

**Affiliations:** Institute for Biochemistry and Molecular Biology, University of Hamburg, Martin-Luther-King-Pl. 6, 20146 Hamburg, Germany; University of Tartu, Institute of Technology, 50411 Tartu, Estonia; Gene Center and Department of Biochemistry, University of Munich, 81377 Munich, Germany

**Author notes:** These authors contributed equally (Agnieszka A. Pochopien, Bertrand Beckert). Daniel N. Wilson. **Email:**.

## Abstract

Protein synthesis is essential to cells and requires a constant supply of nutrients. Amino acid starvation leads to accumulation of uncharged tRNAs that promote ribosomal stalling, which is sensed by the protein kinase Gcn2, together with its effector proteins, Gcn1 and Gcn20. Activation of Gcn2 phosphorylates eIF2, leading to a global repression of translation. Fine-tuning of this adaptive response is performed by the Rbg2/Gir2 complex, which is a negative regulator of Gcn2. Despite the wealth of biochemical data, structures of Gcn proteins on the ribosome have remained elusive. Here we present a cryo-electron microscopy structure of the yeast Gcn1 protein in complex with stalled and colliding 80S ribosomes. Gcn1 interacts with both 80S ribosomes within the disome, such that the Gcn1 HEAT repeats span from the P-stalk region on the colliding ribosome to the A-site region of the lead ribosome. The lead ribosome is stalled in a non-rotated state with peptidyl-tRNA in the A-site, uncharged tRNA in the P-site, eIF5A in the E-site, as well as Rbg2/Gir2 located in the A-site factor binding region. By contrast, the colliding ribosome adopts a rotated state with peptidyl-tRNA in a hybrid A/P-site, uncharged-tRNA in the P/E-site and Mbf1 bound adjacent to the mRNA entry channel on the 40S subunit. Collectively, our findings provide a structural basis for Rbg2/Gir2 repression of Gcn2, and also reveal that colliding disomes are the substrate for Gcn1 binding, which has important implications not only for Gcn2-activated stress responses, but also for general ribosome quality control (RQC) pathways.

## Introduction

All living cells must adapt to a variety of different environmental stresses in a rapid and efficient way to survive. In eukaryotes, the Gcn2 (general control nonderepressible-2) kinase modulates the response to nutrient deprivation by phosphorylation of eukaryotic initiation factor eIF2 on serine 51 of the alpha subunit (Castilho et al., 2014; Hinnebusch, 2005). Although phosphorylation of eIF2 causes a global repression of translation initiation, translation of specific mRNAs also become up-regulated, such as the transcriptional regulator Gcn4 (yeast) or ATF4 (mammals). This in turn induces expression of genes, such as those involved in amino acid biosynthesis, to counteract the amino acid deficiency. The prevailing model for Gcn2 activation during nutrient deprivation is that Gcn2 recognizes and binds ribosomes that have become stalled during translation due to the accumulation of uncharged (deacylated) tRNAs binding to the ribosomal A-site (Castilho et al., 2014; Hinnebusch, 2005). The activation of Gcn2 strictly requires its co-activator Gcn1 (Marton et al., 1993), a large protein (2,672 amino acids or 297 kDa in yeast) conserved from yeast to humans (Castilho et al., 2014). Based on secondary structure predictions Gcn1 is composed almost entirely of HEAT repeats (**Fig. 1a**). The N-terminal three-quarters (residues 1-2052) of Gcn1 are required for tight association with ribosomes *in vivo* (Sattlegger and Hinnebusch, 2000) (**Fig. 1a**) and a reduction in ribosome binding of Gcn1 leads to a concomitant loss in Gcn2 activation (Sattlegger and Hinnebusch, 2005). The central region of Gcn1 is highly homologous to the N-terminal HEAT repeat region of the eukaryotic elongation factor 3 (eEF3) (Marton et al., 1993) (**Fig. 1a**) and overexpression of eEF3 represses Gcn2 activity, suggesting that Gcn1 and eEF3 have overlapping binding sites on the ribosome (Visweswaraiah et al., 2012). The eEF3-like region of Gcn1 is also important for interaction with the N-terminus of Gcn20 (Marton et al., 1997; Vazquez de Aldana et al., 1995) (**Fig. 1a**), a non-essential ATP-binding cassette (ABC) protein that it required for full Gcn2 activity (Vazquez de Aldana et al., 1995). Gcn20 itself does not interact with the ribosome, nor with Gcn2, suggesting that Gcn20 exerts its stimulatory effect on Gcn2 via interaction and stabilization of Gcn1 on the ribosome (Vazquez de Aldana et al., 1995). By contrast, a region (residues 2052-2428) within the C-terminus of Gcn1 mediates direct interaction with the N-terminal RWD domain of Gcn2 (**Fig. 1a**) (Kubota et al., 2001; Kubota et al., 2000; Sattlegger and Hinnebusch, 2000). Moreover, mutations within these regions of either Gcn1 (F2291L or R2259A) or Gcn2 (Y74A) disrupt the Gcn1-Gcn2 interaction, resulting in loss of both eIF2 phosphorylation and derepression of Gcn4 translation (Kubota et al., 2001; Sattlegger and Hinnebusch, 2000). Like Gcn2, Gir2 (Genetically interacts with ribosomal genes 2) contains an N-terminal RWD-domain and interacts with Gcn1 (Wout et al., 2009). Overexpression of Gir2 prevents Gcn2 activation by competing with Gcn2 for Gcn1 binding (Wout et al., 2009). Gir2 forms a complex with the ribosome binding GTPase 2 (Rbg2) (Daugeron et al., 2011; Ishikawa et al., 2013), which has been proposed to dampen the Gcn2 response (Ishikawa et al., 2013). Despite the high conservation and importance of the Gcn pathway, as well as decades of research into the Gcn proteins (Castilho et al., 2014; Hinnebusch, 2005), a structural basis for their mechanism of action on the ribosome has been lacking.

**Figure 1.**
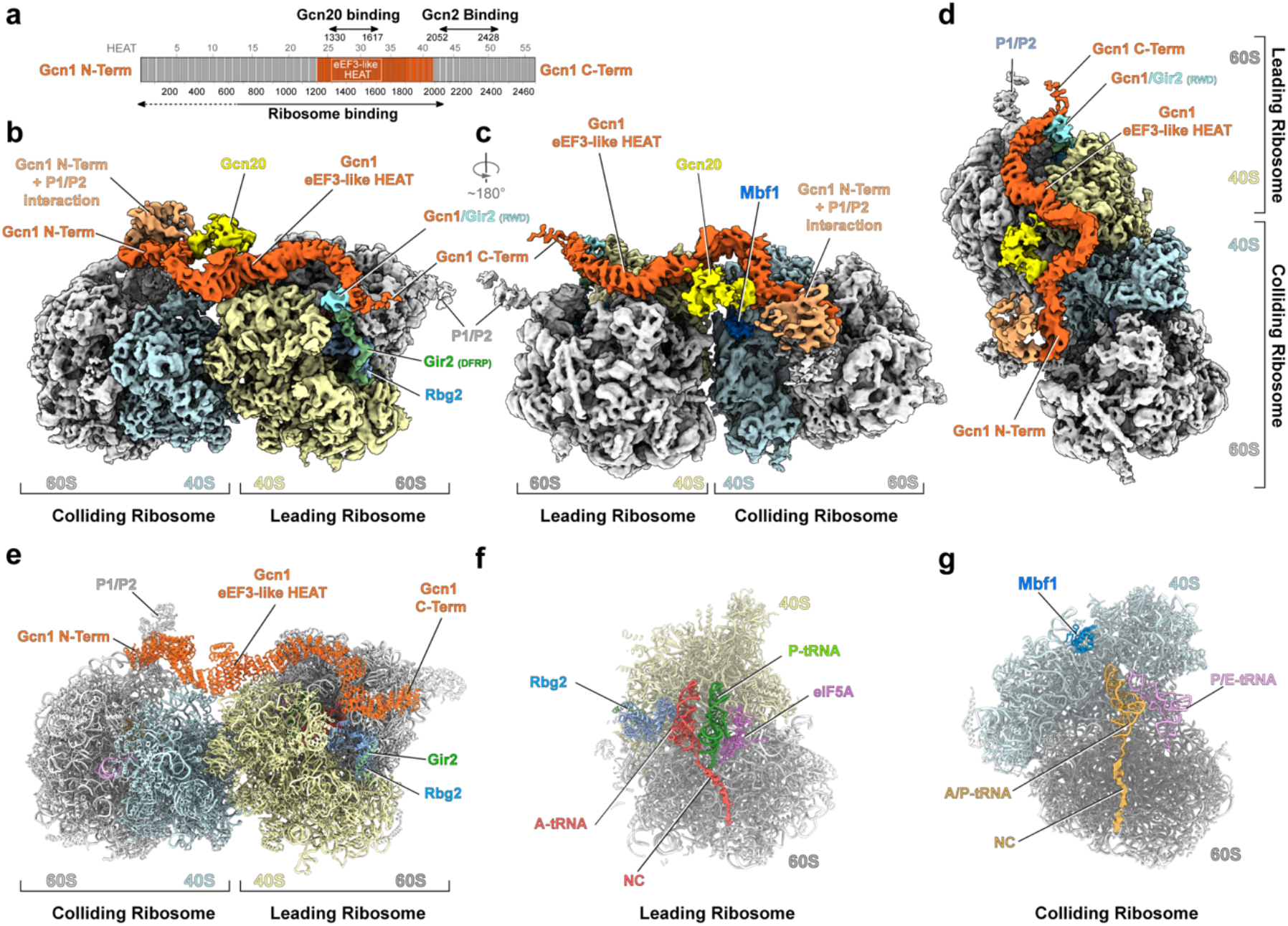
The structure of the Gcn1-bound disome. (**a)** Schematic representation of the yeast Gcn1 protein with its predicted HEAT repeats (grey boxes). Areas of Gcn1 binding to the ribosome, Gcn20 and Gcn2 are indicated with arrows. (**b-d**) Cryo-EM reconstruction of the Gcn1-disome complex with segmented densities for Gcn1 (orange) and Gcn20 (yellow). On the leading ribosome, 40S (cyan), 60S (grey), Rbg2 (light blue), Gir2 (green), Gir2(RWD) (cyan) and on the colliding ribosome, 40S (pale yellow), 60S (grey), Mbf1 (deep blue) and the Gcn1 interaction with P1/P2-stalk proteins (salmon). (**e-g**) Molecular models of the (**e**) Gcn1-bound disome structure, and (**f**) a cut-through view of the Gcn1-disome leading ribosome with Rbg2, peptidyl-tRNA (red) in the A-site, deacylated tRNA (green) in the P-site, eIF5A (purple) in the E-site, and (**g**) the colliding ribosome with Mbf1 (dark blue), peptidyl-tRNA (gold) in the A/P-site and deacylated tRNA (purple) in the P/E-site.

## Results

### Cryo-EM structure of a native Gcn1-disome complex

To investigate how Gcn proteins interact with the ribosome, we set out to determine a cryo-EM structure of a Gcn-ribosome complex. In order to obtain such a complex, we employed affinity chromatography in combination with *S. cerevisiae* cells expressing chromosomally TAP-tagged Gcn20. A C-terminal tag was favoured since the N-terminus of Gcn20 is required for Gcn1 interaction (Marton et al., 1997; Vazquez de Aldana et al., 1995) and C-terminally tagged Gcn20 was previously shown to be indistinguishable from wildtype in complementing □*gcn20* deletion strains (Vazquez de Aldana et al., 1995). Gcn1 is reported to interact with translating ribosomes both in the presence and absence of amino acid starvation (Marton et al., 1997), therefore, purification was performed from mid-log phase cells. Co-purification of Gcn1 and ribosomal proteins with Gcn20-TAP was observed and validated by mass spectrometry (**Dataset S1**). The Gcn20-TAP eluate underwent a mild glutaraldehyde crosslinking treatment before being applied to cryo-grids and subjected to single particle cryo-EM. A low resolution cryo-EM reconstruction of the Gcn20-TAP sample revealed that a minor (5%) subpopulation of ribosomes contained an additional tube-like density, which we assigned to Gcn1 (**Fig. S1**). Interestingly, this class also contained extra density located on the solvent side of the small 40S subunit, which after refinement (to 21 Å) using a larger box-size, revealed that Gcn1 was associated with a disome (two 80S ribosomes), rather than a single 80S monosome (**Fig. S1**).

In order to improve the resolution of the Gcn1-disome complex, we collected 16,823 micrographs on a Titan Krios TEM with a Falcon II direct electron detector. Following 2D classification, the remaining 616,079 ribosomal particles were subjected to 3D classification and divided into 15 different classes (**Fig. S2**). A diverse range of ribosome functional states were identified that did not contain Gcn1, most of which are likely to have co-purified with the polysomes to which the Gcn1-Gcn20-disome was bound. Since many of the states have been previously reported, they will not be discussed further, with the exception of class 5, which contained a post-translocational (P- and E-site tRNAs) state ribosome with eRF1 and eEF3 present (**Fig. S2**). eEF3 has been previously reported to facilitate E-site tRNA release during elongation (Triana-Alonso et al., 1995), however, our results suggest that eEF3 may also perform an analogous function during translation termination. Moreover, since the eEF3 binding site overlaps with Gcn1, and the previous eEF3-ribosome structure was at 9.9 Å (Andersen et al., 2006), we refined the eRF1-eEF3-ribosome structure to an average resolution of 4.2 Å. Of the 15 classes, two classes (1 and 2) contained density that we attributed to Gcn1 (**Fig. S2**) based on overlap with the eEF3 binding site on the ribosome as well as the tube-like density feature characteristic of linear solenoid HEAT repeat proteins (Andrade et al., 2001).

### Gcn1 interact with both the leading stalled and colliding ribosomes

Through further sub-sorting and local refinement (**Fig. S2**), we could obtain a cryo-EM structure of the complete Gcn1-disome with an average resolution of 4.0 Å for the leading stalled ribosome, however, the colliding ribosome was poorly resolved (8.4 Å), indicating some flexibility with respect to the leading ribosome (**Fig. S3a-c**). Thus, we implemented focussed refinement of the individual leading and colliding ribosomes, yielding average resolutions of 3.9 Å and 4.4 Å, respectively (**Fig. S3d-k** and **Table S1**). These maps were combined to generate a cryo-EM map of the complete disome, revealing how density for Gcn1 snakes its way along the disome and fuses at each end with density for the P-stalk proteins of both ribosomes (**Fig. 1b-d** and **Movie S1**). We attributed the extra density contacting Gcn1 at the interface between the leading and colliding ribosomes to the N-terminal domain of Gcn20 (**Fig. 1b-d**) since this region of Gcn1 is critical for interaction with the N-terminus of Gcn20 (Marton et al., 1997; Vazquez de Aldana et al., 1995). However, the density is poorly resolved and therefore no model could be built for this region. In addition to Gcn1, we observed density for Rbg2/Gir2 in the A-site of the leading ribosome as well as multiprotein bridging factor 1 (Mbf1) on the 40S subunit of the colliding ribosome (**Fig. 1b-d** and **Movie S1**), the details and implications of which will be discussed later.

To improve the density for Gcn1, an additional focussed refinement was performed using a mask encompassing Gcn1 and the 40S head of the leading ribosome (**Fig. S3l-o**). The local resolution of Gcn1 was highest (4-7 Å) for the central eEF3-like region of Gcn1 and progressively decreased towards the N- and C-terminal ends (**Fig. S3l-o**). A molecular model for the central region of Gcn1 could be generated based on homology with eEF3 (**Fig. S4a-e**) and individual HEAT repeats could be fitted into the regions flanking the central region (**Fig. 1e** and **Fig. S5a-b**). Analogous to eEF3, the central eEF3-like region of Gcn1 contacts expansion segment 39 (ES39) and ribosomal proteins uS13 and eS19 in the head of the 40S subunit, as well as uL5 and uL18 in the central protuberance of the 60S subunit, of the leading ribosome (**Fig. S5c-d**). The flanking region N-terminal to the eEF3-like region of Gcn1 spans across the disome interface and establishes interactions with eS12 and eS31 within the beak of the 40S subunit of the colliding ribosome (**Fig. S5e-f**). Although we do not observe direct interaction between Gcn1 residues 1060-1777 and eS10, as suggested previously (Lee et al., 2015), we note that eS10 is adjacent to eS12 in the 40S head (**Fig. S5e-f**) and therefore mutations or loss of eS10 could indirectly influence Gcn1 binding to the ribosome. While the central region of Gcn1 is relatively stable, by contrast, the N- and C-terminal “arms” of Gcn1 are highly flexible and winds it way across the disome towards the factor binding site and the P-stalk of the colliding and leading ribosomes (**Fig. 1b-e**). The N-terminal arm of Gcn1 contacts ES43L, uL11 and P0 at the stalk base, whereas the remaining N-terminal 600 aa fuse with density from the other P-stalk proteins, precluding any molecular interpretation (**Fig. S5e-f**). Similarly, the C-terminal arm of Gcn1 also reaches towards the factor binding site, but on the leading ribosome, where the C-terminus appears to extend and contact the P-stalk proteins, although this interaction is also poorly resolved (**Fig. S5g-h**). The interaction between Gcn1 and the P-stalk seen here provides a likely explanation for the observation that mutations or loss of the P-stalk proteins impairs Gcn2-dependent eIF2 phosphorylation (Harding et al., 2019).

The conformation of the leading and colliding ribosomes within the Gcn1-bound disome are distinct from one another. The leading ribosome is in a non-rotated pre-translocational (PRE) state with a peptidyl-tRNA in the A-site, a deacylated tRNA in the P-site and the elongation factor eIF5A in the E-site (**Fig. 1f**). The presence of eIF5A in the Gcn1-disome suggests that translation by the leading ribosome may have slowed down, or even become stalled, due to the presence of problematic polypeptide motifs at the peptidyl-transferase center of the large subunit (Gutierrez et al., 2013; Schuller et al., 2017). By contrast, the colliding ribosome adopts a rotated hybrid state with a peptidyl tRNA in the hybrid A/P-site and a deacylated-tRNA in P/E-site (**Fig. 1g**). In this case, peptide bond formation has ensued but translocation of the mRNA and tRNAs on the small subunit has not occurred. The overall constellation of a non-rotated leading ribosome followed by a rotated colliding ribosome observed in our Gcn1-disome is reminiscent of that observed previously for other colliding disomes, namely, disomes formed in presence of an inactive eRF1^AAG^ mutant (Juszkiewicz et al., 2018) or stalled on CGA-CCG and CGA-CGA containing mRNAs (Ikeuchi et al., 2019; Matsuo et al., 2020), but differs most from the poly(A)-stalled disomes that also contained non-rotated colliding ribosomes (Tesina et al., 2020) (**Fig. S6**).

### Visualization of Rbg2-Gir2 on the leading ribosome of the Gcn1-disome

In addition to the presence of eIF5A in the E-site, the leading ribosome of the Gcn1-disome contained additional density within the factor binding site, adjacent to the A-site, which resembled a GTPase, but not one of the canonical translational GTPases (**Fig. 2a-c**). A mass spectrometry analysis of the Gcn1-disome sample instead revealed the presence of the non-canonical ribosome-binding GTPase 2 (Rbg2), which had comparable intensities to ribosomal proteins as well as some ribosome-associated factors, such as eIF5A and Mbf1 (see **Dataset S1**). Although there is no available structure for Rbg2, it was possible to generate a homology model based on the structure of the closely related (62% identity) Rbg1 (Francis et al., 2012), which could then be satisfactorily fitted to the cryo-EM density (**Fig. S7a-c**). Like Rbg1, Rbg2 comprises four domains; an N-terminal helix-turn-helix (HTH), a C-terminal TGS (ThrRS, GTPase and SpoT) and a central GTPase domain (G-domain) that is interrupted by a ribosomal protein S5 domain 2-like (S5D2L) domain (**Fig. 2d**). In the Gcn1-disome, the TGS domain of Rbg2 interacts with the 40S subunit, contacting helix 5/15 (h5/15) of the 18S rRNA, whereas the G- and HTH domains establish contacts with the 60S subunit, including the stalk base (H43/H44), sarcin-ricin loop (SRL, H95) and H89 (**Fig. 2e-f**). By contrast, the S5D2L domain of Rbg2 makes contacts exclusively with the A-site tRNA, such that α-helix α7 of Rbg2 approaches the minor groove of anticodon-stem in vicinity of nucleotides 27-29 of the A-tRNA (**Fig. 2e-f**). This suggests that Rbg2 can stabilize the accommodated A-site tRNA and thus may work together with eIF5A to facilitate peptide bond formation at problematic peptide motifs. Rbg1 and Rbg2 homologs are encoded in the majority of eukaryotes, with the mammalian counterparts being termed developmentally regulated GTP-binding proteins 1 (DRG1) and DRG2, respectively (Ishikawa et al., 2005). The very high conservation (66% identity) between human (DRG1/2) and yeast (Rbg1/2) orthologs suggests that the interactions observed here for Rbg2 are likely to be identical for DRG2 on the human ribosome.

**Figure 2.**
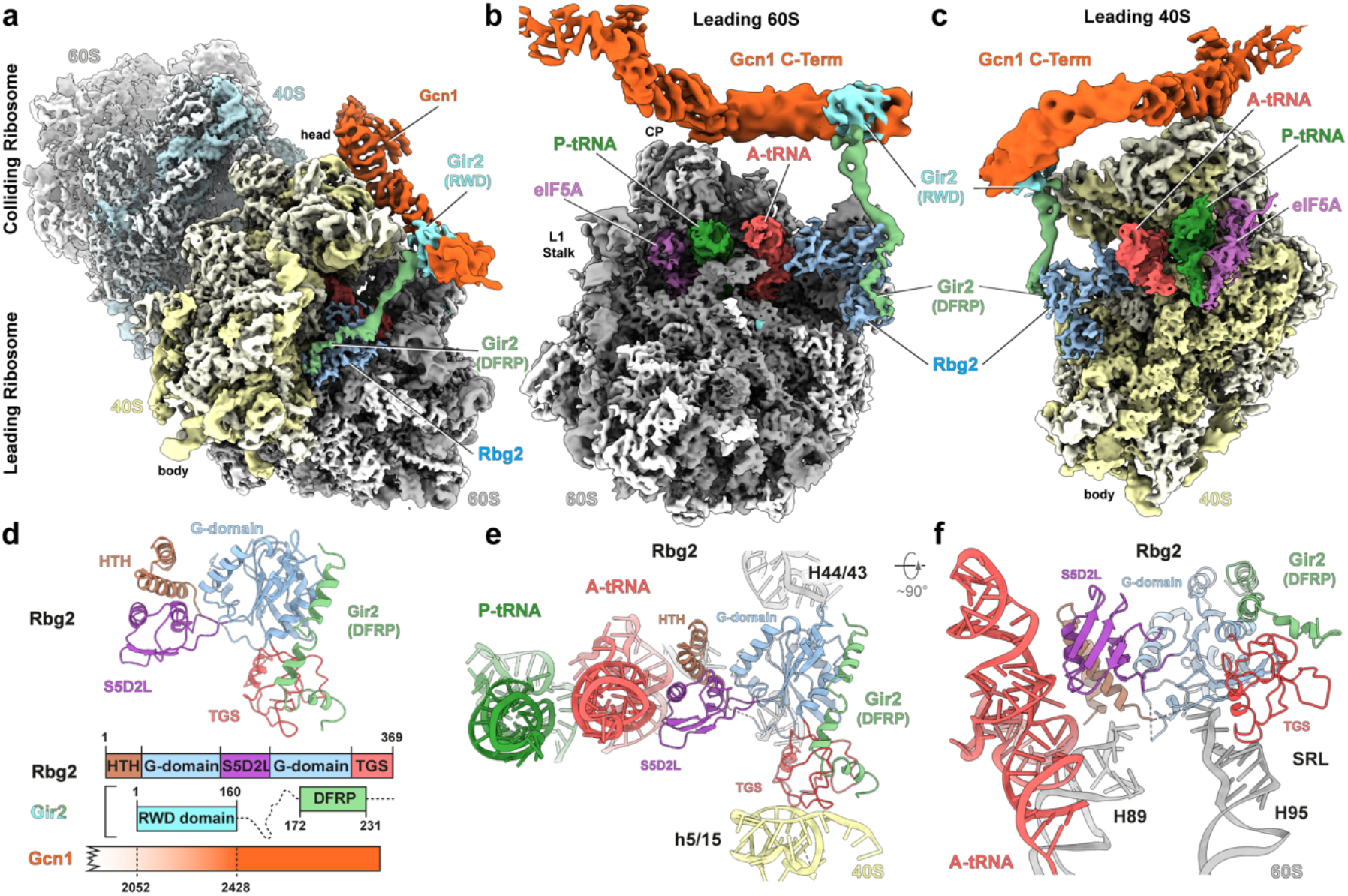
Structure of Rbg2-Gir2 on the leading stalled ribosome. (**a-c**) Cryo-EM reconstruction of the (**a**) Gcn1-disome complex, and interface views of the (**b**) 60S subunit and (**c**) 40S subunit of the leading stalled ribosome. Segmented densities for Gcn1 (orange), colliding ribosome (40S, cyan; 60S, gray), the leading ribosome (40S, pale yellow; 60S, gray), P-tRNA (green), A-tRNA (red), Rbg2 (light blue), Gir2-DFRP (green), Gir2-RWD (cyan) and eIF5A (purple). (**d**) Molecular model for Rbg2-Gir2(DFRP) with schematic representation of the Rbg2-Gir2-Gcn1 interactions. Domains are colored as indicated. (**e,f**) Interactions of Rbg2 coloured by domain as in (**d**) with 40S (pale yellow) and 60S (gray) components and A-site tRNA (red) and P-site tRNA (green).

Under physiological conditions, Rbg2 is very labile, but becomes stabilized through interactions with Gir2, whereas in contrast Rbg1 forms a complex with Tma46 (Ishikawa et al., 2013). Both Gir2 and Tma46 contain a C-terminal DRG family regulatory protein (DFRP) domain that is critical for interaction with Rbg2 and Rbg1, respectively (Daugeron et al., 2011; Ishikawa et al., 2009; Ishikawa et al., 2013). In the Rbg1-Tma46(DFRP) X-ray structure, four α-helices at the C-terminus of the DFRP domain of Tma46 establish contact with the TGS and G-domain of Rbg1 (Francis et al., 2012) (**Fig. S7d**). Consistently, we observe an analogous interaction between the DFRP domain of Gir2 and the TGS and G-domains of Rbg2 (**Fig. 2d** and **Fig. S7e**), however, unlike Tma46 where the linker region wraps around the G-domain of Rbg1, the linker region of Gir2 extends away from Rbg2 towards Gcn1 (**Fig. 2a-c** and **Fig. S7e,f**). This suggests that in the absence of the ribosome, the intimate interaction between Tma46/Gir2 and Rbg1/Rbg2 stabilizes their respective complexes, whereas upon ribosome binding, the N-terminal domains are freed to find new interaction partners. With respect to Rbg2-Gir2 in the Gcn1-disome structure, we observe that the density for the N-terminal region of Gir2 fuses with the C-terminal region (residues 2000-2200) of Gcn1 (**Fig. 2a-c**). Although the contact cannot be resolved in any detail due to the high flexibility within this region, support for such an interaction is well-documented; firstly, the N-terminal RWD domain of Gir2 is necessary and sufficient for interaction with Gcn1 and, secondly, a construct containing only the C-terminal residues 2048-2382 of Gcn1 retains the ability to bind Gir2 (Wout et al., 2009). Moreover, the ribosome-association of Gir2 was also shown to be partially dependent on the presence of Gcn1 (Wout et al., 2009). Thus, taken together with the biochemical studies, our structural findings reveal that Gir2 indeed acts as a physical link between Rbg2 and Gcn1 and is likely to prevent Gcn2 activation in situations where Rbg2 mediates successful restoration of translation of the leading ribosome by stimulating peptide bond formation.

### Visualization of Mbf1 on the colliding ribosome of the Gcn1-disome

Within the colliding ribosome of the Gcn1-disome, we observed additional density located between the head and body of the 40S subunit that we attributed to Mbf1 (**Fig. 3a,b** and **Fig. S8a**), a conserved archaeal/eukaryotic protein that suppresses +1 frameshifting at inhibitory CGA-CGA codon pairs in yeast (Wang et al., 2018). Recent findings indicate that the mammalian Mbf1 homolog, EDF1, stabilizes GIGYF2 at collisions to inhibit translation initiation in cis (Juszkiewicz et al., 2020a; Sinha et al., 2020), however, in our reconstruction, we did not observe any additional density for the yeast GIGYF2 homolog. The assignment of Mbf1 was based on (i) the high intensity of Mbf1 peptides in the mass spectrometry analysis of the Gcn1-disome sample (see **Dataset S1**) and (ii) the excellent agreement between the cryo-EM density and a homology model for Mbf1 generated from the NMR structure of the C-terminal helix-turn-helix (HTH) domain of Mbf1 from the fungus *Trichoderma reesei* (Salinas et al., 2009) (**Fig. S8b,c**). Moreover, the binding site for Mbf1 observed here on the Gcn1-disome is consistent with that observed recently on stalled disomes/trisomes (Sinha et al., 2020). The C-terminal HTH domain of Mbf1 connects h33 in the head with h16 and h18 within the body of the 40S subunit (**Fig. 3c**), and thereby stabilizes a non-swiveled conformation of the head (**Fig. S8d-f**). Interaction with h33 is likely to be critical for Mbf1 function since mutations (I85T, S86P, R89G, R89K) within helix α3, which contacts h33, leads to loss of frameshift suppression (Wang et al., 2018). Binding of Mbf1 leads to a shift of h16 towards the body of the 40S subunit (**Fig. 3d**), which is stabilized by interactions of the C-terminus and helix α6 of Mbf1 with the minor groove of h16 (**Fig. 3d**). In addition to the HTH domain, we were able to model the N-terminal residues 25-79 of Mbf1, including two short α-helices α1 and α2 formed by residues 26-37 and 59-68 (**Fig. 3b**). Helix α2 of Mbf1 interacts directly with helices α1 and α2 of ribosomal protein uS3 (**Fig. 3e**). These interactions are likely to be essential for Mbf1 function since S104Y and G121D substitutions within these two helices of uS3 result in a loss of frameshift suppression, which can be partially restored by overexpression of Mbf1 (Wang et al., 2018). Mutations (R61T and K64E) located in the distal region of helix α2 of Mbf1 also abolish frameshift suppression (Wang et al., 2018). Arg61 comes into hydrogen bonding distance with Asn111 of S3 (**Fig. 3e**) and Lys64 appears to interact with the backbone phosphate oxygens of h16. Asc1 (RACK1) is also critical for suppressing +1 frameshifting at CGA repeats (Wang et al., 2018; Wolf and Grayhack, 2015), however, our structure suggests that this is not due to direct interaction with Mbf1, but rather because Asc1 appears to be critical for disome formation by establishing multiple interface contacts between the leading and colliding ribosome (**Fig. S6**).

**Figure 3.**
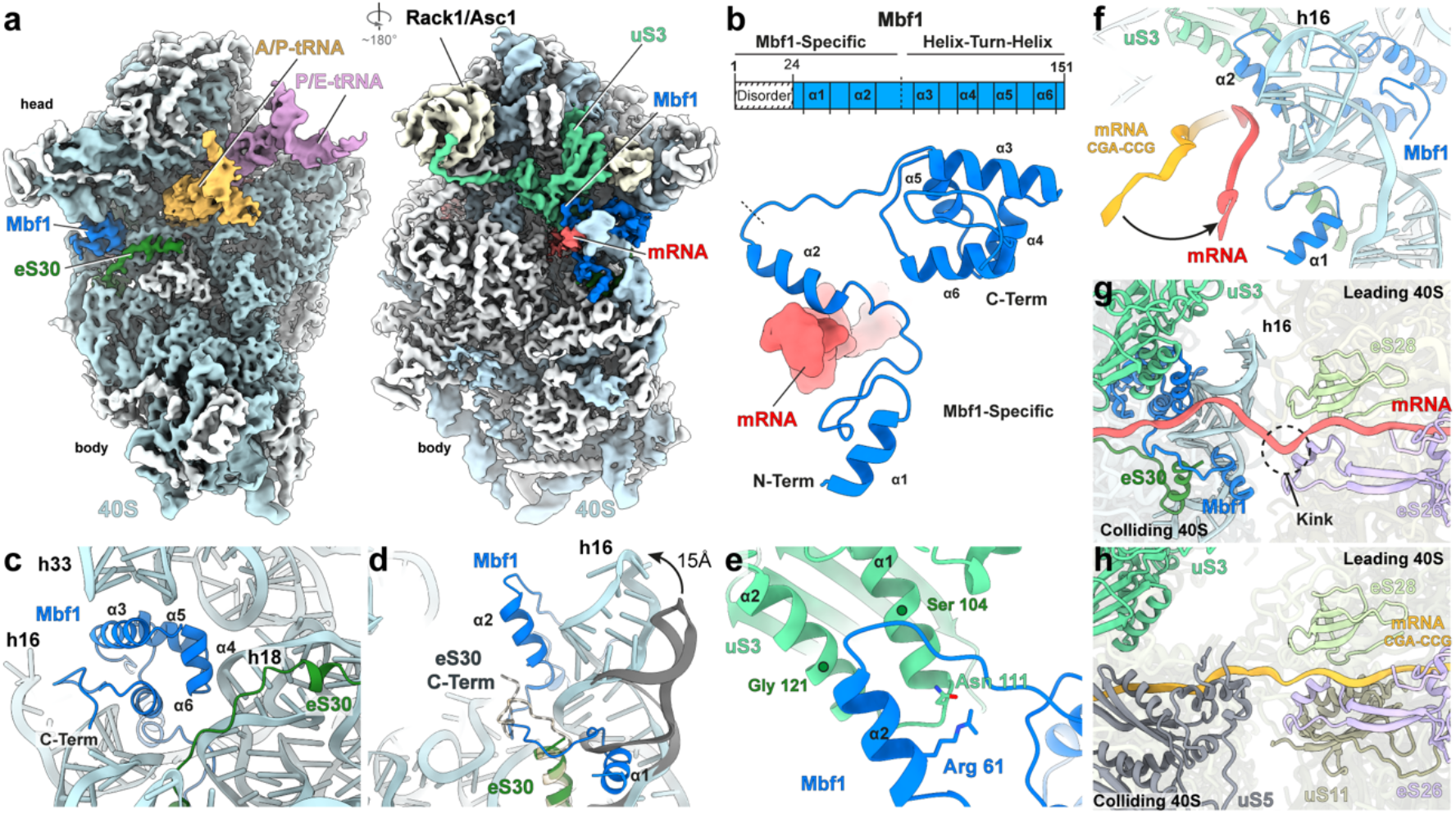
Structure of Mbf1 on the colliding ribosome. (**a**) Cryo-EM map of the interface (left) and solvent (right) view of the 40S subunit from the colliding ribosome, with 40S proteins (white), 18S rRNA (cyan), A/P-tRNA (gold), P/E-tRNA (purple), eS30 (dark green), uS3 (light green), mRNA (red) and Mbf1 (deep blue). (**b**) Schematic and cartoon representation of the Mbf1 molecular model (deep blue). (**c**) Mbf1 helix-turn-helix motif bound to the 40S subunit. (**d**) Refolding of h16 and eS30 C-terminus destabilization upon Mbf1 binding. Comparison of h16 (cyan) and eS30 (dark green) in the Mbf1-bound colliding ribosome to h16 (gray) and eS30 (light gold) in the colliding ribosome of an Mbf1-lacking disome (PDB ID 6SNT) (Matsuo et al., 2020) (**e**) Close-up view on uS3 and Mbf1 helix 2 interactions. (**f**) Comparison of the path of the mRNA (red) in the Mbf1-bound structure compared to the mRNA (orange) in a colliding ribosome of an Mbf1-lacking disome (PDB ID 6I7O) (Ikeuchi et al., 2019). (**g,h**) The mRNA path between the 40S of the colliding and the 40S of the leading ribosome within (**g**) the Gcn1-disome and (**h**) the CGA-CCG stalled disome (PDB ID 6I7O) (Ikeuchi et al., 2019). Components interacting with the mRNA at the 40S-40S interface are shown for each disome, respectively.

The N-terminal residues preceding helix α2 of Mbf1 wrap around the stem of h16, and, as a consequence, the C-terminus of eS30e becomes disordered (**Fig. 3d,f** and **Fig. S8g-i**). Helix α1 of Mbf1 is located within the major groove of h16 oriented towards the interface, suggesting that N-terminal 24aa that are not observed in our structure may reach towards the leading ribosome (**Fig. 3f,g**). The shift of h16 induced by Mbf1 brings the minor groove of h16 into contact with the mRNA, which together with direct interaction observed between the mRNA and helix α2 of Mbf1, causes a redirection in the path of the 3’ end of the mRNA, when compared to the CGA-CGG stalled disome structure (**Fig. 3f**) (Ikeuchi et al., 2019; Matsuo et al., 2020). Moreover, the mRNA appears to be kinked at the interface between the leading and colliding ribosomes, suggesting a relaxed state is adopted instead of a more extended and potentially strained conformation seen in other disomes (**Fig. 3g**). Ribosome collisions have been shown to induce +1 frameshifting because the colliding ribosome exerts a pulling force on the mRNA during translocation that promotes slippage of the mRNA with respect to the tRNAs in the leading ribosome (Simms et al., 2019). Our findings suggest that Mbf1 suppresses +1 frameshifting on the leading ribosome by binding to the colliding ribosome and locking the 40S subunit in such a manner that mRNA movement is prevented. Specifically, Mbf1 prevents head swiveling that is required for mRNA/tRNA translocation and stabilizes the mRNA via direct interactions as well as indirectly by promoting additional interactions between the mRNA and the ribosome, especially h16. Finally, we note that the overall arrangement of the leading and colliding ribosomes in the Gcn1-disome is more compact than observed for the CGA-CCG disome (Ikeuchi et al., 2019; Matsuo et al., 2020) (**Fig. 3g,h** and **Fig. S9a-f**). This results in a shorter path that the mRNA needs to traverse between the ribosomes, which may also contribute to maintaining a relaxed mRNA conformation on the leading ribosome. These findings imply that Gcn1 interaction with the 40S subunit of the colliding ribosome (**Fig. S5e,f**), rather than Mbf1-40S interactions, are likely to be critical for promoting the novel compact architecture of the Gcn1-disome.

## Discussion

Collectively, our findings enable us to present a model for how Gcn1 can sense stalled ribosomes and subsequently recruit Gcn2 to the stalled disome. Under conditions of amino acid starvation, the binding of deacylated tRNA in the A-site of the ribosome causes translational stalling, which in turn increases the frequency of ribosome collisions and disome formation (Darnell et al., 2018; Meydan and Guydosh, 2020) (**Fig. 4a**). We envisage that such disomes are recognized by the Gcn1-Gcn20 complex in an analogous manner to that observed here (**Fig. 4b**), and that Gcn1 can recruit and activate Gcn2 via direct interaction with its N-terminal RWD domain (**Fig. 4c**), analogous to the interaction established between Gcn1 and the RWD domain of Gir2. Moreover, there is growing evidence that Gcn2 activation also occurs in response to stimuli that promote collisions, but in a deacylated tRNA independent manner (Anda et al., 2017; Deng et al., 2002; Ishimura et al., 2016; Wu et al., 2020) (**Fig. 4d**). This is consistent with our structural findings showing that Gcn1 recognizes the architecture of a disome, rather than directly monitoring the presence or absence of an A-site tRNA. Thus, an important conclusion from our study is that recognition of colliding ribosomes or disomes enables Gcn1 to facilitate Gcn2 activation in response to many diverse environmental stresses that act to inhibit translation. Furthermore, the discovery of Rbg2-Gir2 bound to the leading ribosome of the Gcn1-disome supports the concept that Gcn1 acts as a scaffold to interact with other accessory factors and thereby fine-tune the level of Gcn2 activation (Castilho et al., 2014). One such example is the Rbg2-Gir2 complex, which is known to repress Gcn2 activation (Wout et al., 2009). Recent findings suggest that Rbg2 (and Rbg1) facilitate translation through problematic polybasic (Arg/Lys-rich) stretches in proteins (Zeng et al., 2020). Consistently, we observe Rbg2 interacting with the A-site tRNA on the leading ribosome of the Gcn1-disome, suggesting that it may stabilize the A-tRNA to promote peptide bond formation and restore the translational activity of the stalled ribosome (**Fig. 4e,f**). Moreover, during this “problem-solving” phase, we suggest that Gcn2 recruitment and activation is prevented due to the competing interaction of the RWD domain of Gir2 with Gcn1 (Wout et al., 2009) (**Fig. 4e**). Thus, a second important conclusion from our study is that Gcn1-disome provides a platform upon which factors can bind to regulate Gcn2 activation and in the case of Rbg2-Gir2, the repression of Gcn2 is coupled with the potential reactivation of translation on the leading ribosome.

**Figure 4.**
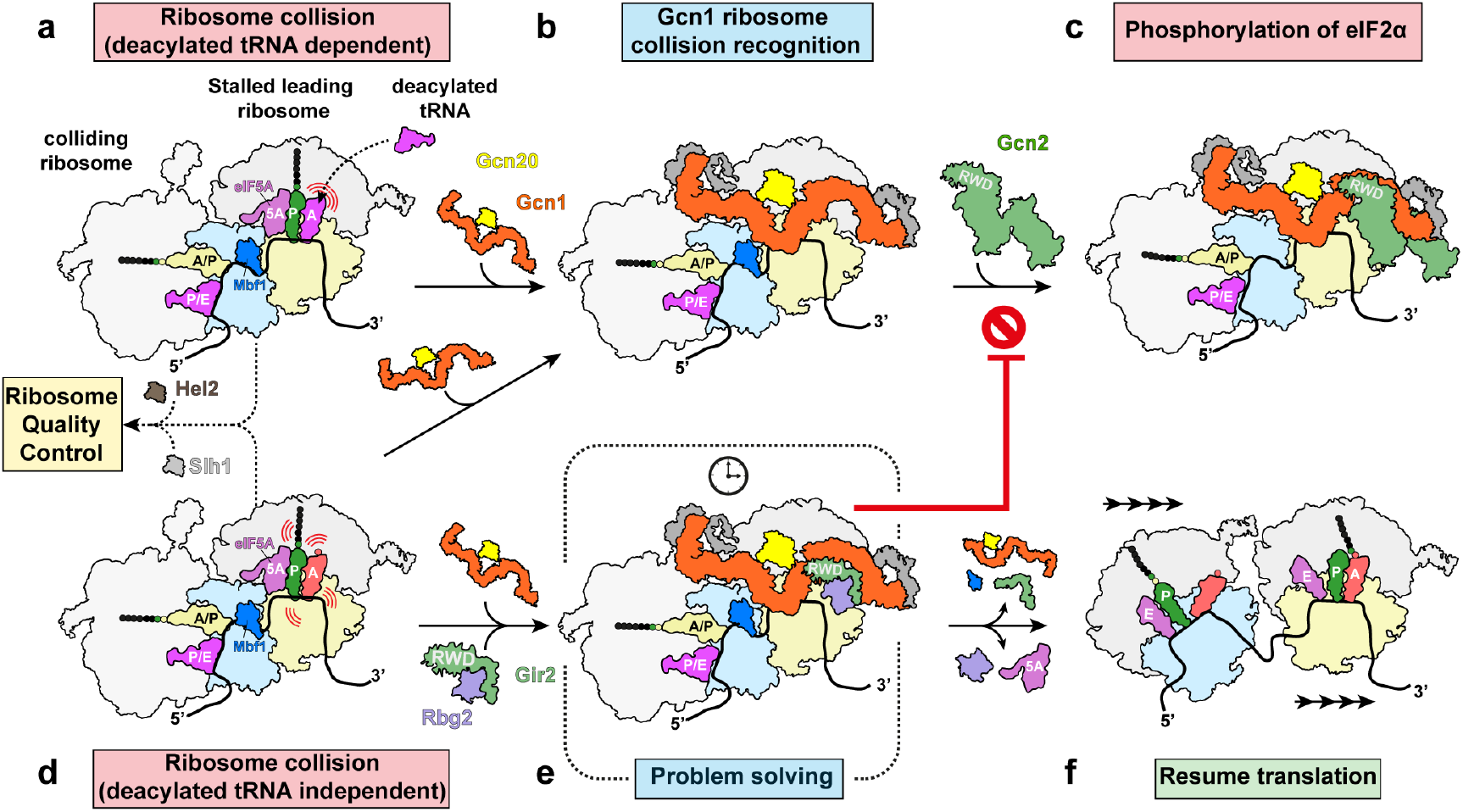
Gcn1 as a checkpoint for disomes collision. (**a**) Amino acid starvation leads to an increased binding of uncharged tRNAs within the ribosomal A-site leading to translation slow-down/stalling and collisions. (**b**) Colliding ribosomes (disomes) are recognized by Gcn1-Gcn20. (**c**) Gcn1 in turn recruits Gcn2 via direct interaction with Gcn2 N-terminal RWD domain. Activation of Gcn2 results in the phosphorylation of eIF2α and induction of the GAAC pathway. (**d**) Translating ribosomes may also encounter specific mRNA sequences/structures and/or nascent polypeptide motifs that induce a translational slow-down or pausing, also leading to collisions and disome formation, which are known as substrates for the ribosome quality control, but are also recognized by Gcn1. (**e**) Concomitant recruitment of the Rbg2-Gir2 to the leading ribosome by Gcn1 allows Rbg2 to resolve the slow down, while Gir2 prevents the recruitment and activation of Gcn2, and eventually (**f**) allows translation to resume.

Finally, our finding that colliding ribosomes are the substrate for Gcn1 recruitment provides a rationale for the emerging link between Gcn2 activation and the ribosome quality control (RQC) pathway. The RQC pathway targets stalled ribosomes for disassembly and promotes degradation of aberrant mRNAs and nascent polypeptide chains (Brandman and Hegde, 2016; Inada, 2020; Joazeiro, 2019). A central player in the RQC pathway is Hel2/ZNF598, which recognizes colliding ribosomes and ubiquitylates specific 40S ribosomal proteins. This in turn recruits the helicase Slh1/ASCC to dissociate the leading ribosome from the mRNA, allowing the following ribosomes to continue translating (Juszkiewicz et al., 2020b; Matsuo et al., 2020). Deletion of Hel2 in yeast has been recently reported to cause an increase in eIF2 phosphorylation (Meydan and Guydosh, 2020). In light of our results, a likely explanation for this observation is that in the absence of Hel2, additional disome substrates become available for Gcn1 binding, leading to increased Gcn2 activation and eIF2 phosphorylation. Like Hel2, Slh1 is also non-essential in yeast, but loss of Slh1 is synthetic lethal when combined with a Rbg1/Tma46/Rbg2/Gir2 quadruple knockout (Francis et al., 2012). This implies that for survival, eukaryotic cells must remove the translational roadblock by either reactivation of translation of the leading ribosome by for example Rbg2-Gir2 (**Fig. 4e,f**) or disassembly of the stalled disome roadblock on the mRNA via the RQC pathway. While further work will be required to dissect out the mechanistic details and interplay between the factors and pathways, their conservation across all eukaryotes, including humans, implies a central and evolutionary importance.

## Acknowledgments

We thank Charlotte Ungewickell for preparation of cryo-grids. This research was supported by grants from the Deutsche Forschungsgemeinschaft WI3285/8-1 and SPP-1879 (to D.N.W.), the European Union from the European Regional Development Fund through the Centre of Excellence in Molecular Cell Engineering (2014-2020.4.01.15-0013 to T.T.); and the Estonian Research Council (PRG335 to T.T.).

## Supplementary Information

### Cell extract extraction and sucrose gradient analysis

Yeast whole-cell extracts of TAP-tagged GCN20 strain (SC0000; MATa; ura3-52; leu2-3,112; YFR009w::TAP-KlURA3) Euroscarf, were prepared from cultures grown to mid log-phase (OD_600_ of 0.8-1.0) in either YPD or SC medium (plus supplements as required). Cells were harvested by centrifugation at 4,400 x g for 10 min at 4°C in a Sorvall SLC-6000 rotor (Marshall Scientific). The cell pellets were washed with ice cold water and lysis buffer (20 mM HEPES (pH 7.4), 100 mM KOAc, 10 mM Mg(OAc)_2_, 1 mM DTT, Complete EDTA-free Protease Inhibitor cocktail (Roche)), transferred to a 50 ml falcon, resuspended in lysis buffer and disrupted using glass beads. Glass beads were removed by centrifugation for 5 min at 4,400 x g at 4°C in the Rotanta 460R falcon centrifuge (Hettich). The cell debris were further pelleted by centrifugation at 17,600 x g for 15 min at 4°C in a Sorvall RC 6 SS-34 rotor.

### Tandem affinity purification for cryo-EM analysis

The TAP-Tag *in vivo* pull-out was performed using the GCN20 TAP-tagged strain essentially as described before (Schmidt et al., 2016b). Cells were harvested at the mid log phase at an OD_600_ of 2.5 and lysed via glass bead disruption. The cleared lysate was incubated with IgG-coated magnetic DynabeadsR©M-270 Epoxy (Invitrogen) for 1 h at 4°C with slow tilt rotation. The elution was performed by addition of AcTEV Protease (Invitrogen) for 2 h at 17°C in elution buffer containing 20 mM HEPES (pH 7.4). 100 mM KOAc, 10 mM Mg(OAc)_2_, 1 mM DTT and 1 mM ADPNP (Sigma-Aldrich). The pull-out sample was analyzed by 4-12% SDS-PAGE and immunoblot analysis.

### Proteomics sample preparation and nano-LC/MS/MS analysis

Gcn20-pullout sample was loaded onto SDS-PAGE gels and run for a short time so that the samples entered into the gel and then the complete sample was excised from the gel as a single band. The gel band was then destained in 1:1 acetonitrile (ACN):100 mM ammonium bicarbonate (ABC) with vortexing, reduced with 10 mM dithiothreitol at 56°C and alkylated with 50 mM chloroacetamide in the dark. Protein digestion was carried out overnight with 10 ng/μl of dimethylated porcine trypsin (Sigma Aldrich) in 100 mM ABC at 37°C. Peptides were extracted from the gel matrix using bath sonication, followed by 30 min vortexing in 2 volumes of 1:2 5% formic acid (FA): ACN. The organic phase was evaporated in a vacuum-centrifuge, after which the peptides were desalted on in-house made C18 (3M) solid phase extraction tips. Purified peptides were reconstituted in 0.5% trifluoroacetic acid. Peptides were injected to an Ultimate 3000 RSLCnano system (Dionex) using a 0.3 × 5 mm trap-column (5 μm C18 particles, Dionex) and an in-house packed (3 μm C18 particles, Dr Maisch) analytical 50 cm × 75 μm emitter-column (New Objective). Peptides were eluted at 200 nL/min with an 8-40% (30 min) A to B gradient (buffer A: 0.1% (v/v) FA; buffer B: 80% (v/v) ACN + 0.1% (v/v) FA) to a quadrupole-orbitrap Q Exactive Plus (Thermo Fisher Scientific) MS/MS via a nano-electrospray source (positive mode, spray voltage of 2.5 kV). The MS was operated with a top-10 data-dependent acquisition strategy. Briefly, one 350-1,400 m/z MS scan at a resolution setting of R = 70,000 was followed by higher-energy collisional dissociation fragmentation (normalized collision energy of 26) of the 10 most intense ions (z: +2 to +6) at R = 17,500. MS and MS/MS ion target values were 3,000,000 and 50,000 ions with 50 and 100 ms injection times, respectively. Dynamic exclusion was limited to 15 s. MS raw files were processed with the MaxQuant software package (version 1.6.1.0) (Tyanova et al., 2016). Methionine oxidation, protein N-terminal acetylation were set as potential variable modifications, while cysteine carbamidomethylation was defined as a fixed modification. Identification was performed against the UniProt (www.uniprot.org) *Saccharomyces cerevisiae* (strain ATCC 204508 / S288c) reference proteome database using the tryptic digestion rule. Only identifications with at least 1 peptide ≥ 7 amino acids long (with up to 2 missed cleavages) were accepted. Intensity-based absolute quantification (iBAQ) (Schwanhausser et al., 2011) feature of MaxQuant was enabled. This normalizes protein intensities by the number of theoretically observable peptides and enables rough intra-sample estimation of protein abundance. Peptide-spectrum match, peptide and protein false discovery rate was kept below 1% using a target-decoy approach (Elias and Gygi, 2007). All other parameters were default. Data are available via ProteomeXchange with identifier PXD021365.

### Sample and grid preparation

0.02% glutaraldehyde were added to the freshly eluted TAP-Tag pull-out complex and incubated for 20 min on ice. The crosslinking reaction was quenched by addition of 25 mM Tris-HCl (pH 7.5) and n-dodecyl-D-maltoside (DDM) was added to a final concentration of 0.01% (v/v). 5 μL (8 A_260_/mL) of the freshly purified and crosslinked complex was applied to 2 nm precoated Quantifoil R3/3 holey carbon supported grids and vitrified using a Vitrobot Mark IV (FEI, Netherlands).

### Low resolution data collection and image processing

The TAP-Tag *in vivo* Gcn20 pull-out sample was initially checked by generating a low resolution cryo-EM reconstruction with a dataset consisting of 264 micrographs collected on a 120 kV Tecnai G2 Spirit (FEI) transmission electron microscope (TEM) equipped with a TemCam-F816 camera (TVIPS) at a pixel size of 2.55 Å with a defocus range of −3.5 to −1.5 μm. Particle picking was performed automatically using Gautomatch (http://www.mrc-lmb.cam.ac.uk/kzhang/) resulting in 40,012 particles (**Fig. S1**). An initial 3D reconstruction was obtained using a vacant *S. cerevisiae* 80S ribosome as a reference, followed by 3D classification into 5 classes. When compared to the vacant 80S ribosome, class 1 (5%; 2,033 particles) displayed additional density within the A-site factor binding site as well as a tube-like extra density spanning the head and central protuberance of the 80S ribosome. In addition, the class 1 ribosome displayed clear density for a neighbouring ribosome, suggesting the presence of a disome. Class 2 (14%; 5,589 particles) revealed a vacant 80S ribosome whereas class 3 (17%; 6,809 particles) had some density within the A-site region. Finally, class 4 and 5 were attributed to “junk” classes. The particles from class 1 were selected and re-extracted with bigger box size (increased from 420 to 700) in order to encompass the full neighbouring ribosome and subsequently 3D refined at around 21 Å resolution, revealing the structure of disome with the extra “worm-like” density for Gcn1 spanning both ribosomes within the disome (**Fig. S1**).

### High resolution cryo-EM data collection and image processing

To obtain high resolution of the Gcn1-disome complex, a total of 16,823 micrographs with a total dose of 25 e^−^/Å^2^ at a nominal pixel size of 1.084 Å and with a defocus ranging from −2.8 to −1.3 μm were collected using the EPU software (Thermo Fisher) on a FEI Titan Krios TEM (Thermo Fisher) operating at 300 kV equipped with a Falcon II direct electron detector. Each micrograph, consisting of a series of 10 frames, was summed and corrected for drift and beam-induced motion using MotionCor2 (Zheng et al, 2017). The power spectra, defocus values, astigmatism and estimation of micrograph resolution were determined using Gctf (Zhang, 2016). Automated particle picking was then performed using Gautomatch (http://www.mrc-lmb.cam.ac.uk/kzhang/), yielding initially 949,522 particles that were subjected to 2D classification using the RELION-3.0 software package (Zivanov et al., 2018) (**Fig. S2**). After 2D classification, 616,079 particles were selected and subsequently subjected to 3D refinement using a vacant *S. cerevisiae* 80S ribosome as initial reference. The initially refined particles were further 3D classified into 15 classes (**Fig. S2**). Class 1 (8.6%, 52,933 particles) was identified as the leading stalled ribosome containing densities for A- and P-site tRNAs, eIF5A, Gcn1, Rbg2 and Gir2, whereas class 2 (6.5%, 39,575 particles) was identified as the collided ribosome with densities for Gcn1, Mbf1, A/P- and P/E-site tRNAs. Class 3 (24.2%, 149,887 particles) and class 4 (24.4%, 151,167 particles) represent the major classes, both containing tRNAs. Class 3 had rotated ribosomes with hybrid A/P- and P/E-site tRNAs, analogous to that observed previously (Behrmann et al., 2015; Svidritskiy et al., 2014), whereas class 4 was non-rotated with A- and P-site tRNAs. In addition, class 4 had additional density in the A-site that would be consistent with an open conformation of EF-1a that occurs after tRNA release and GTP hydrolysis (Pittman et al., 2006). Class 5 (10.6%, 65,173 particles) contained non-rotated ribosomes with P- and E-site tRNAs and additional density for eRF1 in the A-site and eEF3 bound to the head and central protuberance of the 40S and 60S, respectively. Further sub-sorting of this class revealed subpopulations containing eEF3 and eRF1, eRF1 and eIF5A, as well as eRF1, substoichiometric eRF3 and eIF5A, however, these subpopulations could not be refined to high resolution due to the low particle number. Class 6 (7.7%, 47,171 particles) contained hibernating 80S ribosomes with the presence of eEF2 and Stm1, as reported previously (Anger et al., 2013; Brown et al., 2018). Class 7 (1.3%, 8,169 particles) contained mature 60S particles, whereas the remaining classes (classes 8-15 totalling 16.7%, 102,004 particles) contained damaged and/or non-aligning particles are were considered as low resolution/junk.

In order to ensure that Gcn1-containing particles were not lost during the initial 3D classification, all classes (1-15) were pooled together and sub-sorted again but using a mask to focus sorting on Gcn1. This resulting in a major class with 88,453 particles. Sub-sorting of this class produced a high-resolution class (class 1 with 30,016 particles) as well as two low resolution or orientation biased classes (classes 2 and 3, totally 58,437 particles). The particles from class 1 were selected and re-extracted with bigger box size (increased from 420 to 700) in order to encompass the full neighbouring ribosome. Following 3D refinement, CTF refinement yielded a structure of the full disome (**Fig. S2**) with an average resolution of 4.0 Å for the leading stalled and 8.4 Å for the colliding ribosome (**Fig. S3a-c**). Local refinement of the of the individual ribosomes yielded average resolutions of 3.9 Å (**Fig. S3d-g**) and 4.4 Å, respectively (**Fig. S3h-k**). Local refinement was also performed on the 40S head/Gcn1 region of the leading stalled ribosome (**Fig. S2**), which improved the local resolution of Gcn1 (**Fig. S3l-o**). Finally sharpening of the final maps was performed by dividing the maps by the modulation transfer function of the detector and by applying an automatically determined negative B factor to the maps using Relion-3.0. For model building the final maps were locally filtered and the local resolution estimated using Relion-3.0. The final resolution of each volume was determined using the “gold standard” criterion (FSC = 0.143).

### Molecular Modelling

A homology model of the Gcn1 eEF3-like HEAT repeat region (predicted residue 1330-1641) was created using the 80S-bound eEF3 model (this study) as a template for SWISS-MODELLER (Bienert et al., 2017). Comparison of the two cryo-EM maps of the ribosome bound by eEF3 and Gcn1 revealed an identical overall conformation and ribosomal binding site of eEF3-HEATs and Gcn1 eEF3-like HEATs. Based on that similarity, the created Gcn1 model (8 HEAT repeats; residues 1324-1638) was fitted into the appropriate 7 Å low-pass filtered cryo-EM map using the command “fit in map” in UCSF Chimera 1.13.1 (Pettersen et al., 2004) and manually adjusted with Coot version 0.8.9.2 (Emsley and Cowtan, 2004). Beyond the Gcn1 EF3-like HEAT repeat region, three poly-alanine HEAT repeats were *de novo* modelled in the N-terminal direction (residues 1216-1323) and eight C-terminal HEAT repeats (residues 1639-1922) using Coot version 0.8.9.2 (Emsley and Cowtan, 2004). The peripheral N- and C-terminal regions could not be modelled due high flexibility of these regions in the cryo-EM map, however, the tube-like features of the density is consistent with secondary structure predictions of additional HEAT repeats, therefore, we tentatively fitted HEAT repeats from residues 600 to 2600. The very N- and C-terminal regions fuse with the stalk proteins and could not be modelled. Although Gcn20 was not well resolved, the location of the density and interaction region with Gcn1 would be consistent with the N-terminal region of Gcn20.

The extra density at the A-site entry factor of the stalled-leading ribosome was identified and modelled as Rbg2-Gir2. The crystal structure of the Rbg1 protein (PDB ID 4A9A) (Francis et al., 2012) was rigid body fitted into the isolated density using Chimera and the structure of Rgb2 was then generated by homology modelling within Coot (Emsley and Cowtan, 2004) and Isolde (Croll, 2018) and subsequently refined in Phenix (Adams et al., 2010). Gir2-DFRP model was obtained using a homology model based on the Tma46 template from PDB ID 4A9A. However, due to the lack of side chain information, Gir2-DFRP has been modelled only as poly-Ala. Finally, to generate a full molecular model for the stalled leading Gcn1-Rbg2-Gir2-80S complex, existing models for the translating *S. cerevisiae* ribosome (PDB ID 6I7O), Phe-tRNA for A-tRNA and P-tRNA (PDB ID 1EVV), eIF5A (PDB ID 5GAK) (Schmidt et al., 2016a) were used and combined with the Rbg2-Gir2 and Gcn1 model. The molecular model was then refined using Phenix (Adams et al., 2010).

The extra density located between the head and body of the colliding ribosome was identified and modelled as Mbf1 protein. The NMR structure of the C-terminal part of Mbf1 protein from the fungus *Trichoderma reesei* (PDB ID 2JVL) was rigid body fitted into the isolated density using Chimera and the *S. cerevisiae* structure of Mbf1 C-terminal region (residues 80-140) was then modelled by homology in Coot, while the N-terminal region (residues 24-79) was *de novo* modelled using Coot and Isolde. Helix 1 of Mbf1 was placed in the extra-density located in the major groove of h16, however, due to the lack of resolution, the model is only tentative and consists of a polyalanine trace. To generate a full molecular model for the colliding Gcn1-Mbf1-80S complex, existing models for the translating *S. cerevisiae* ribosome PDB ID 6I7O), Phe-tRNA (PDB ID 1EVV) were used and combined with the Mbf1 and Gcn1 model. The molecular model was then refined using Phenix.

### Figure preparation

Figures showing atomic models and electron densities were generated using either UCSF Chimera (Pettersen et al., 2004) or Chimera X (Goddard et al., 2018) and assembled with Inkscape (https://inkscape.org/) and Adobe Illustrator.

**Fig. S1.**
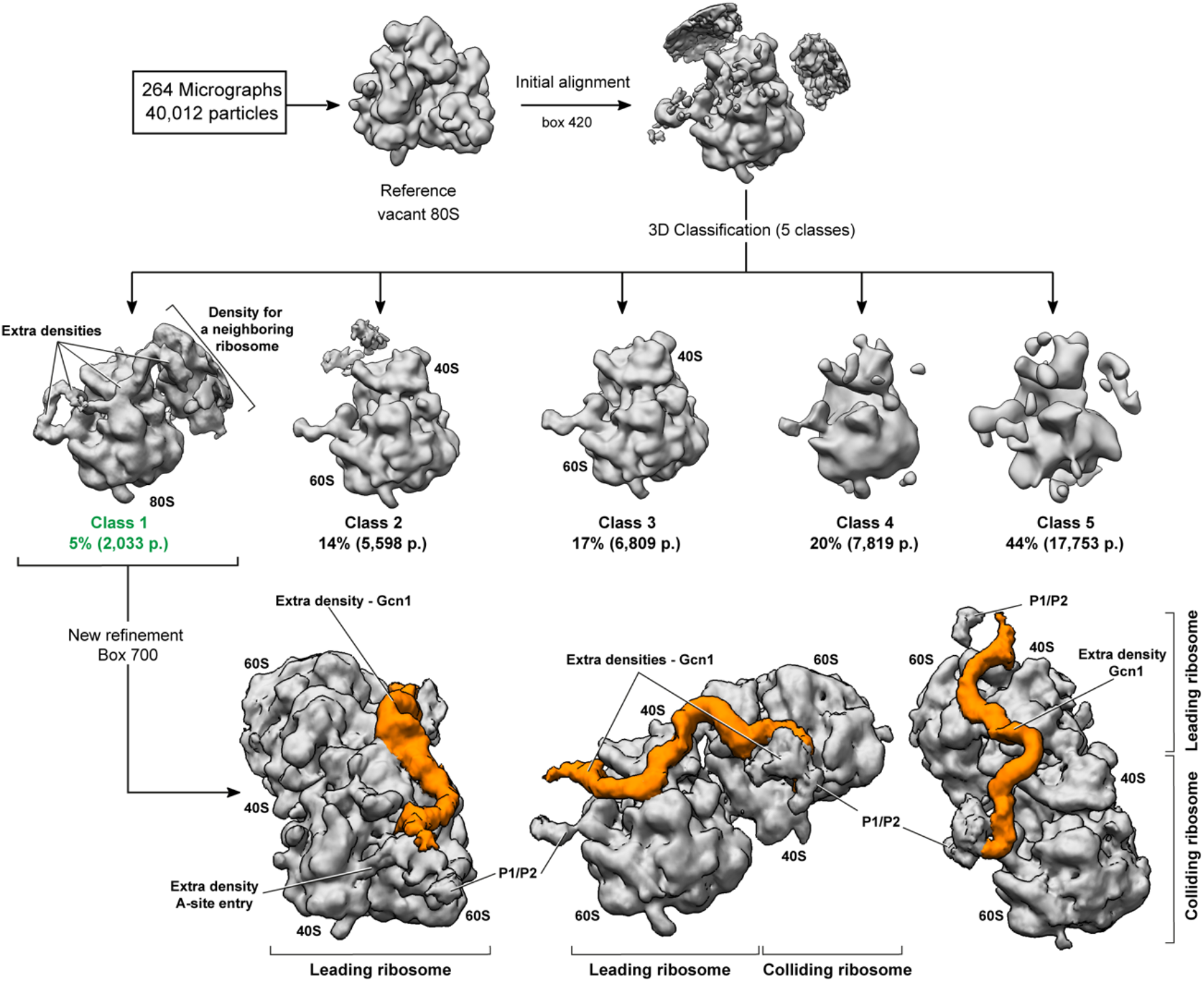
Processing of the low resolution Gcn20-TAP dataset. After automatic acquisition of 264 micrographs, 40,012 particles were selected for the initial reconstruction with an initial alignment using a vacant *S. cerevisiae* 80S ribosome. The particles were then subjected to 3D classification, sorting the particles into 5 classes. Class 2 and 3 contain vacant ribosomes without any extra density at the nominal resolution of 21 Å. Class 1 with the extra densities was then refined using a bigger box size and shows a disome with an extra density connecting the two ribosomes (orange) as well as an extra density at the A-site entry.

**Fig. S2.**
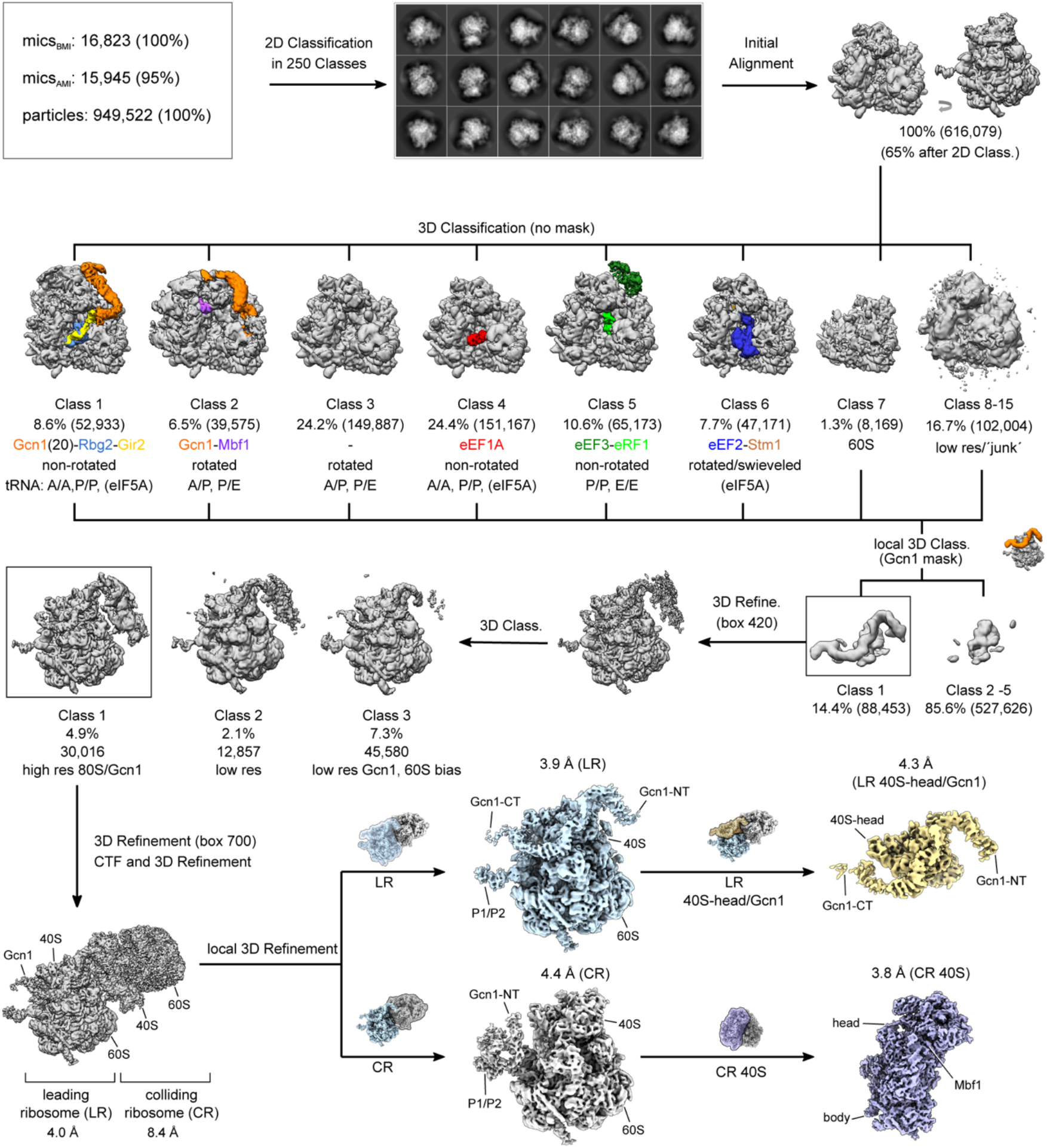
Processing of the high resolution Gcn20-TAP dataset. After manual inspection (AMI) (before manual inspection, BMI) of 16,823 micrographs, 15,945 were selected for the initial particle picking, resulting in 949,522 particles. Following 2D classification, 616,079 particles were initially aligned against a vacant *S. cerevisiae* 80S ribosome and subjected to 3D classification, sorting the particles into 15 initial classes. Class 1 (8.6%, 52,933 particles) revealed a density for Gcn1-Rbg2-Gir2 bound leading stalled ribosome whereas class 2 (6.5%, 39,575 particles) was identified as the colliding ribosome bound by Mbf1. Class 3 (24.2%, 149,887 particles) showed a ribosomal species in a rotated state bearing hybrid A/P and P/E-tRNAs. In the two subsequent classes a non-rotated 80S ribosome was identified bound by A/A- and P/P-tRNA, eIF5A and eEF1A (class 4, 24.4%, 151,167 particles) or P/P- and E/E-tRNA, eEF3 and eRF1 (class 5, 10.6%, 65,173 particles). Class 6 (7.7%, 47,171 particles) represented a rotated ribosome with swivelled 40S head occupied by eIF5A, eEF2 and Stm1. Class 7 (1.3%, 8,169 particles) represented a 60S whereas the rest of the classes (16.7%, classes 8-15) were identified as low-resolution species. All classes (1-15) were combined and also subjected to local sorting using a mask for Gcn1. Class 1 (14.3%, 88,453 particles) containing Gcn1 bound particles was subsequently refined and 3D classified into three classes. High resolution particles in Class1 (4.9%, 30,016 particles) were again 3D and CTF refined with an enlarged box size and resulted in a final disome reconstruction at 4.0 Å for the leading and 8.4 Å for the colliding ribosome. The leading (blue) and the colliding ribosome (grey) were masked and individually local refined resulting in maps with a final resolution at 3.9 Å and 4.4 Å, respectively. To increase the resolution of Gcn1 and Mbf1, the leading and the colliding ribosome was subjected to an additional local refinement with an individual mask encompassing the important regions: the 40S head and Gcn1 (yellow mask) for the leading ribosome and 40S (purple mask) for the colliding ribosome, which resulted in final reconstructions of the masked regions at a final resolution of 4.3 Å and 3.8 Å, respectively.

**Fig. S3.**
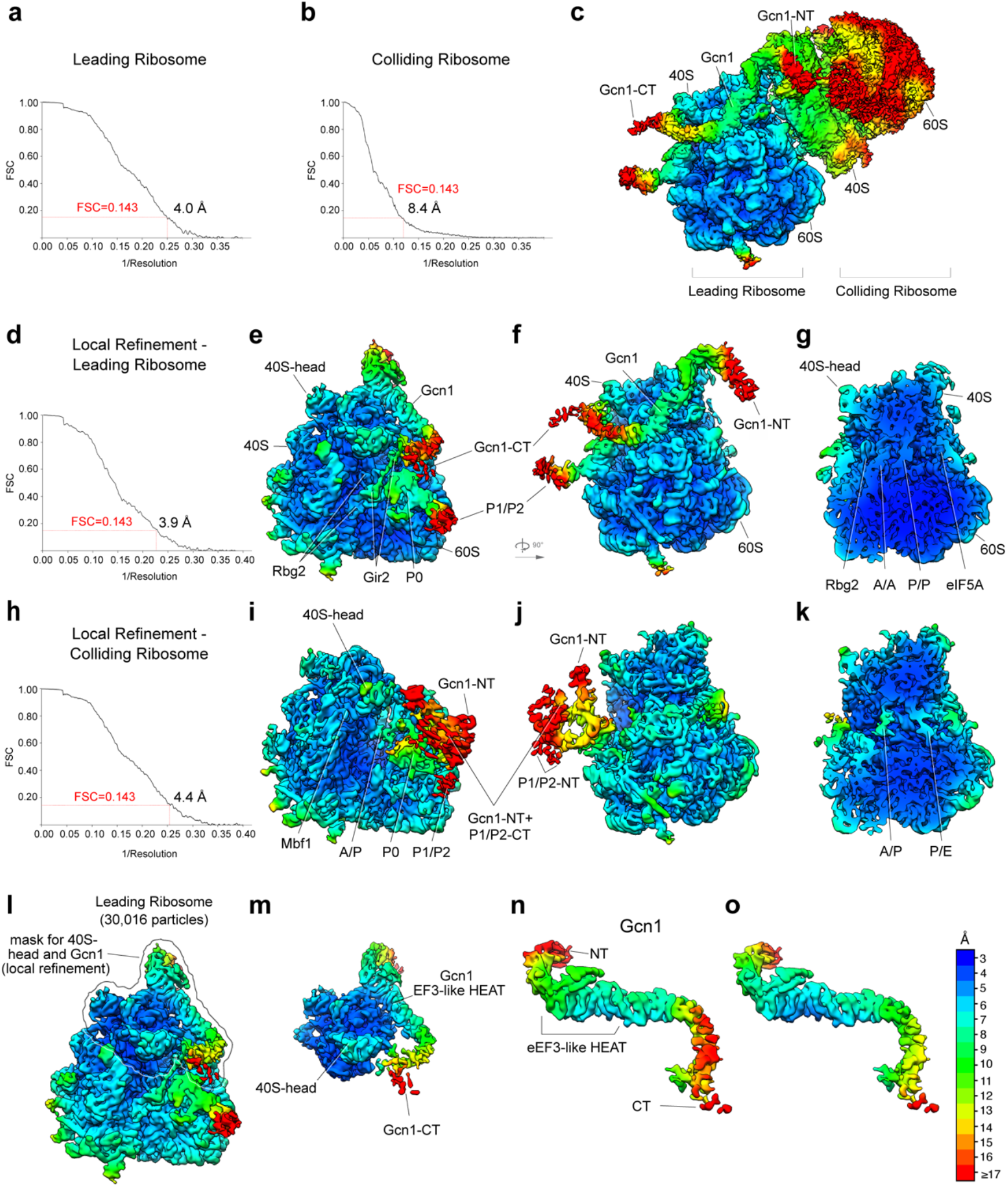
Local resolution of cryo-EM reconstructions of the Gcn1-disome. (**a-c**) Cryo-EM reconstruction of the Gcn1-disome, with (**a-b**) Fourier shell correlation (FSC) curve of the final reconstructions indicating an average resolution of 4.0 Å for the leading and 8.4 Å for the colliding ribosome, according to the gold-standard criterion (FSC=0.143). (**c**) Cryo-EM map of the full disome filtered to 8 Å and colored according to local resolution. (**d-g**) Cryo-EM reconstruction of the leading ribosome, with (**d**) FSC curve of the final reconstruction indicating an average resolution of 3.9 Å (FSC=0.143). (**e,f**) Cryo-EM reconstruction of the leading ribosome filtered to 8 Å and colored according to the local resolution. (**g**) Transverse section of the volume shown in (**f**). (**h-k**) Cryo-EM reconstruction of the colliding ribosome, with (**h**) FSC curve of the final reconstruction indicating an average resolution of 4.4 Å (FSC=0.143). (**i,j**) Cryo-EM reconstruction of the leading ribosome filtered to 9 Å and colored according to the local resolution. (**k**) Transverse section of the volume shown in (**j**). (**l**) Cryo-EM reconstruction of the locally refined leading ribosome colored according to local resolution. The border represents the mask, which was used for the local refinement of Gcn1 including Gcn1 and the head of the 40S subunit of the leading ribosome. (**m**) Postprocessed volume shown in (**l**). (**n-o**) Local resolution of Gcn1 (**n**) before and (**o**) after the local refinement.

**Fig. S4.**
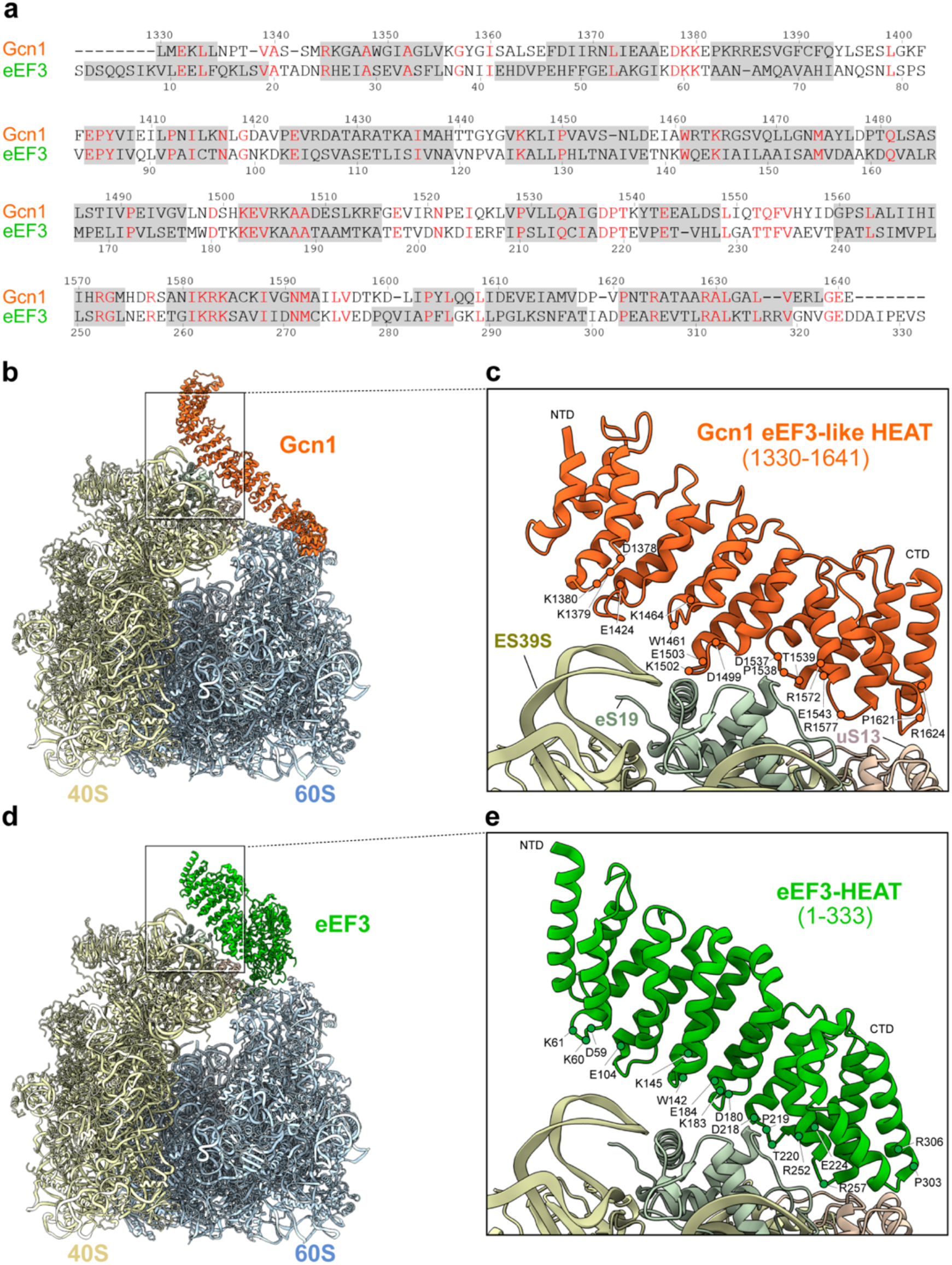
Comparison between the Gcn1 eEF3-like HEAT region and eEF3 HEAT repeats. (a) Sequence alignment of the eEF3-like HEAT repeat region of Gcn1 with HEAT repeats in eEF3. Conserved residues are denoted in red and helices are marked as grey boxes. (b) Molecular model of the Gcn1-80S complex (Gcn1, orange; 40S, pale yellow; 60S, cyan) with (c) zoomed view showing the detailed amino acid composition of the Gcn1 region interacting with ES39S (pale yellow), eS19 (pale green), uS13 (pale coral) of the 40S on the leading ribosome. (d) The eEF3 ribosomal model (eEF3, green) with (e) enlarged view of the eEF3 interface contacting the ribosome

**Fig. S5.**
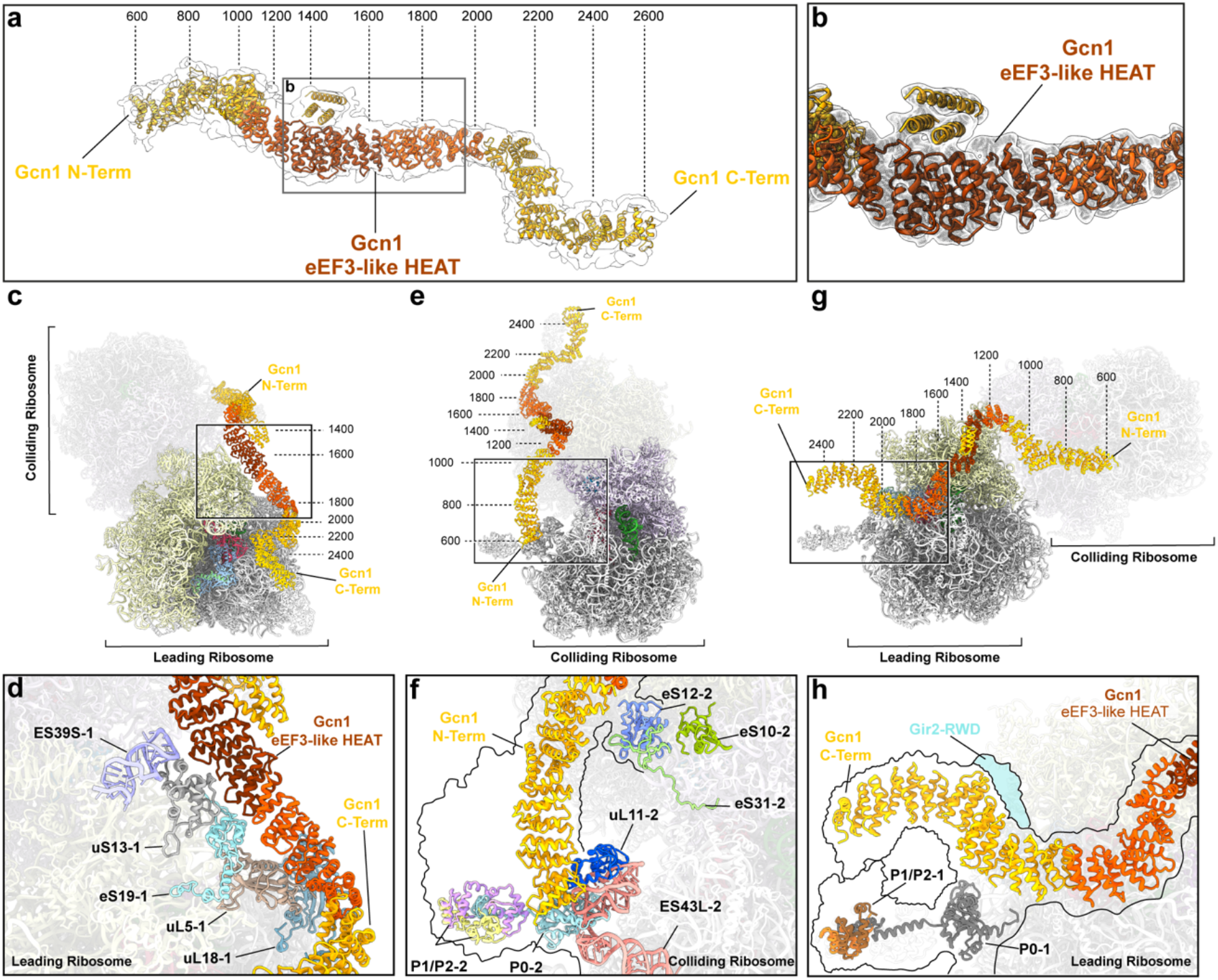
Interactions of Gcn1 within the disome. (**a**) Molecular model of the Gcn1 within the cryo-EM density (gray transparent). The eEF3-like region (dark orange) is shown enlarged in (**b**). The bright orange HEAT repeats were fitted individually based on the features of the density for helices, whereas the yellow HEAT repeats are tentative placements fitted to the map to obtain an approximate estimation of the HEAT positions within Gcn1. (**c-h**) Interactions of Gcn1 within the disome, specifically focussed on the (**c-d**) central region of Gcn1 interacting with ES39S, uS13 and eS19 of the 40S head and uL5 and uL18 of the central protuberance of the leading ribosome, (**e-f**) N-terminal region of Gcn1 interacting with uL11, ES43L and the P-stalk proteins P0, P1 and P2 of the colliding ribosome, and (**g-h**) the C-terminal region of Gcn1 interacting with the RWD domain of Gir2 (cyan) as well as the P-stalk proteins P0, P1 and P2 of the leading ribosome. The border shown in (**f**) and (**h**) reflects the density within the cryo-EM map.

**Fig. S6.**
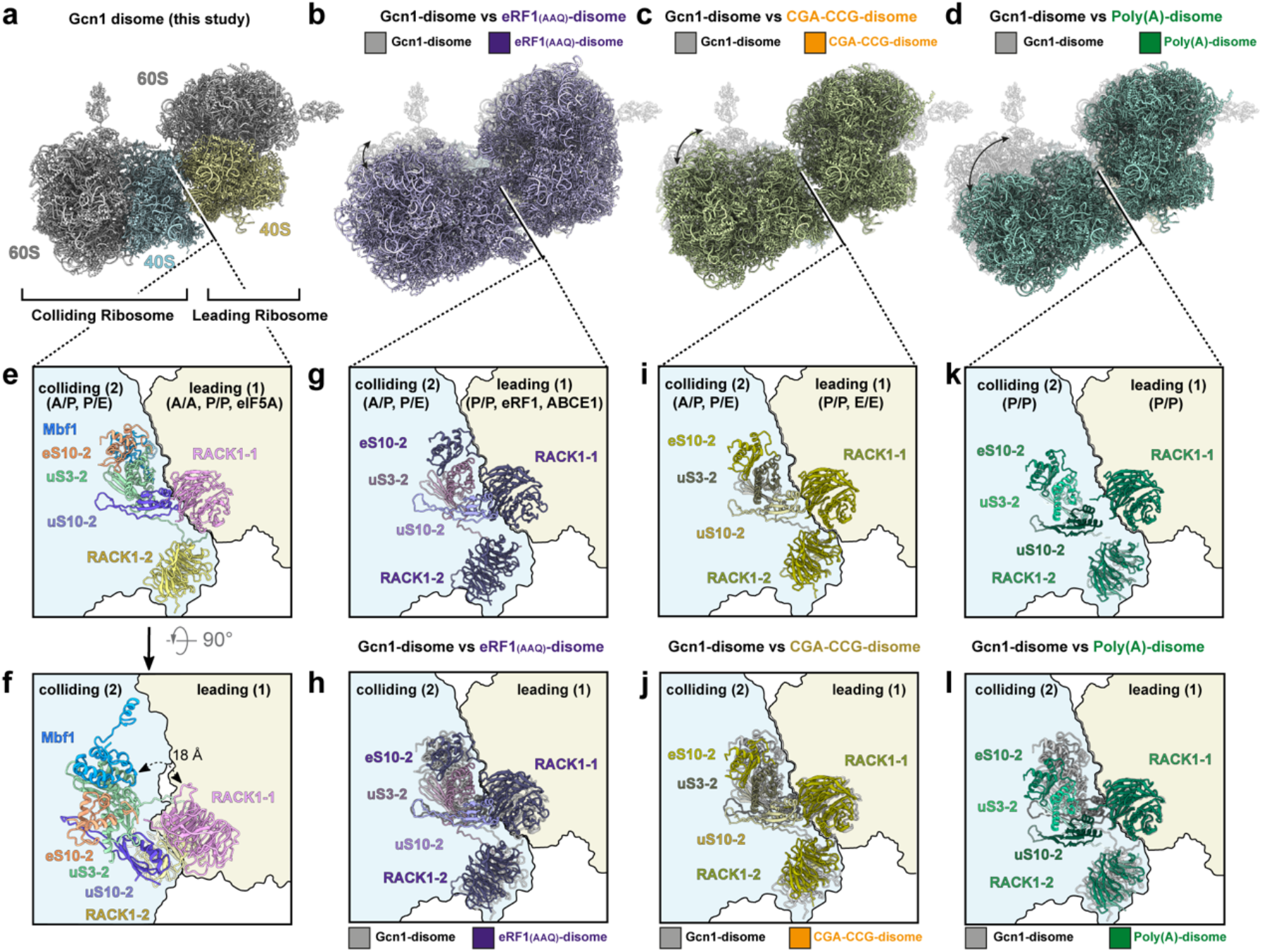
40S-40S interaction interface of the Gcn1-bound disome and its comparison to other disomes. (**a**) Overview of the Gcn1-disome molecular model. (**b**) Collided disome stalled by a catalytically nonactive eRF1 mutant (eRF1-AAQ) (PDB ID: 6hcm, 6hcq) (Juszkiewicz et al., 2018) (purple) overlaid with the Gcn1-disome (grey) shown in (**a**). (**c**) Disome stalled on a CGA-CCG mRNA (PDB ID: 6i7o) (Ikeuchi et al., 2019) (orange) overlaid with the Gcn1-disome (grey) structure shown in (**a**). (**d**) Collided disome stalled on poly(A) mRNA (PDB ID: 6t83) (Tesina et al., 2020) (dark green) overlaid with the Gcn1-disome (grey) structure shown in (**a**). (**e,f**) Close-up view on the Gcn1-disome interaction interface of the 40S subunits from the leading stalled ribosome (pale yellow) and the colliding ribosome (pale blue). (**g**) Close-up view on the interaction interface from the collided disome stalled by a eRF1-AAQ mutant. (**h**) Overlay of (**e**) and (**g**). (**i**) Close-up view on the interaction interface from the disome stalled on a CGA-CCG mRNA. (**j**) Overlay of (**e**) and (**i**). (**k**) Close-up view on the interaction interface from the collided disome stalled on poly(A) mRNA. (l) Overlay of (**e**) and (**k**).

**Fig. S7.**
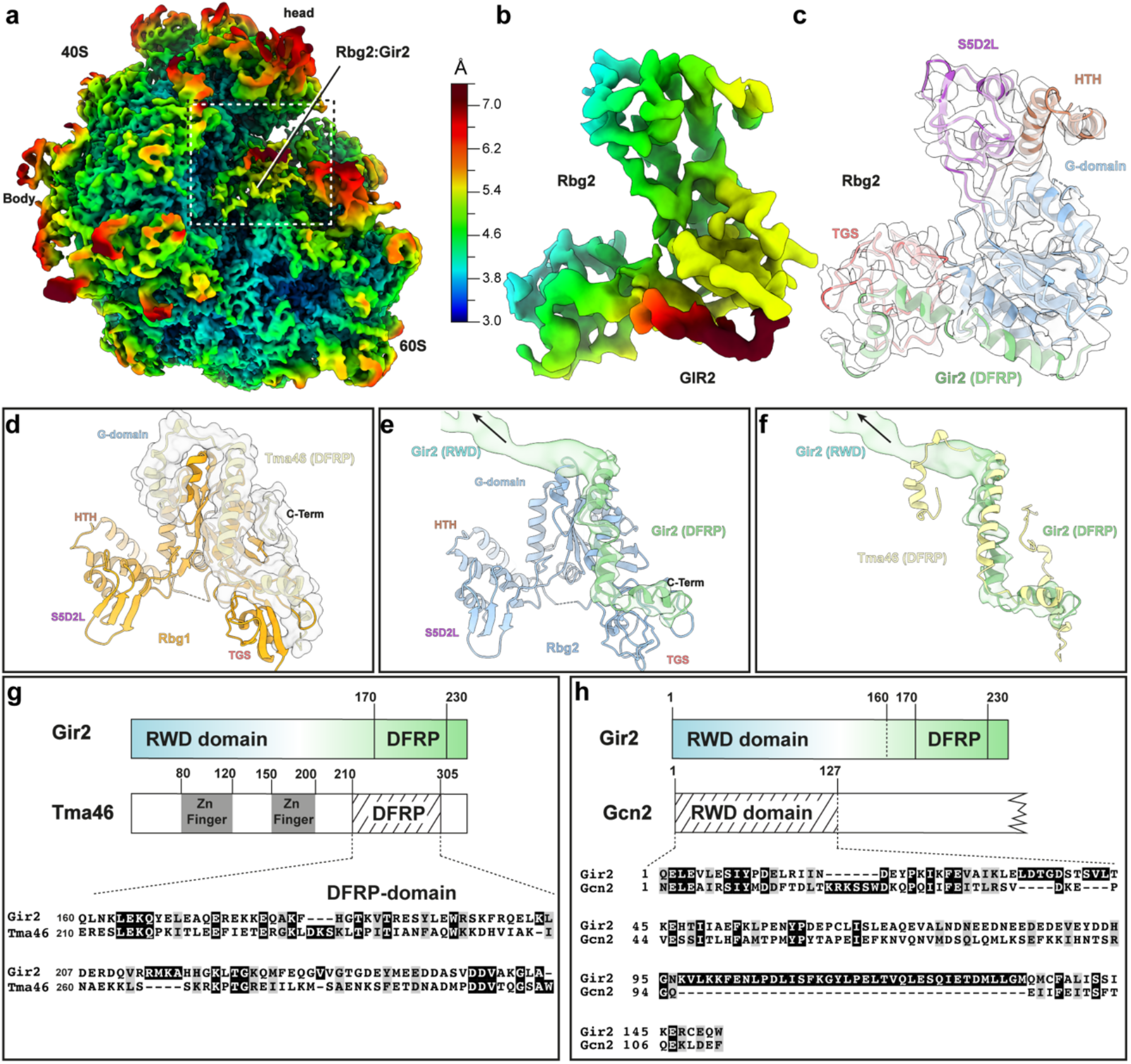
Molecular model for Rbg2 and comparison of Gir2 with Tma46 and Gcn2. (**a**) Local resolution map of the leading ribosome with the binding site of Rbg2 indicated. (**b**) Isolated density of the Rbg2-Gir2 complex from the Gcn1-disome, coloured according to local resolution. (**c**) Molecular model for Rbg2 fitted into isolated cryo-EM density for Rbg2 (transparent grey) and colored due to domain organization. (**d**) Structure of Rbg1 (orange, PDB ID: 4A9A) (Francis et al., 2012) in complex with Tma46(DFRP) domain (light yellow with transparent surface). (**e**) Conformation of Rbg2 (blue) and Gir2(DFRP) domain (green with transparent surface) as found bound to a stalled leading ribosome in the Gcn1-disome. (**f**) comparison of cryo-EM density and model for Gir2 (green) with the model for Tma46 as obtained by aligning the Rbg1-Tma46 complex to Rbg2. (**g**) Schematic representation of the domain structure of Gir2 and Tma46 with zoom on a sequence alignment of their respective DFRP domains. (**h**) Schematic representation of the domain structure of Gir2 and Gcn2 with a zoom on a sequence alignment their respective RWD domains.

**Fig. S8.**
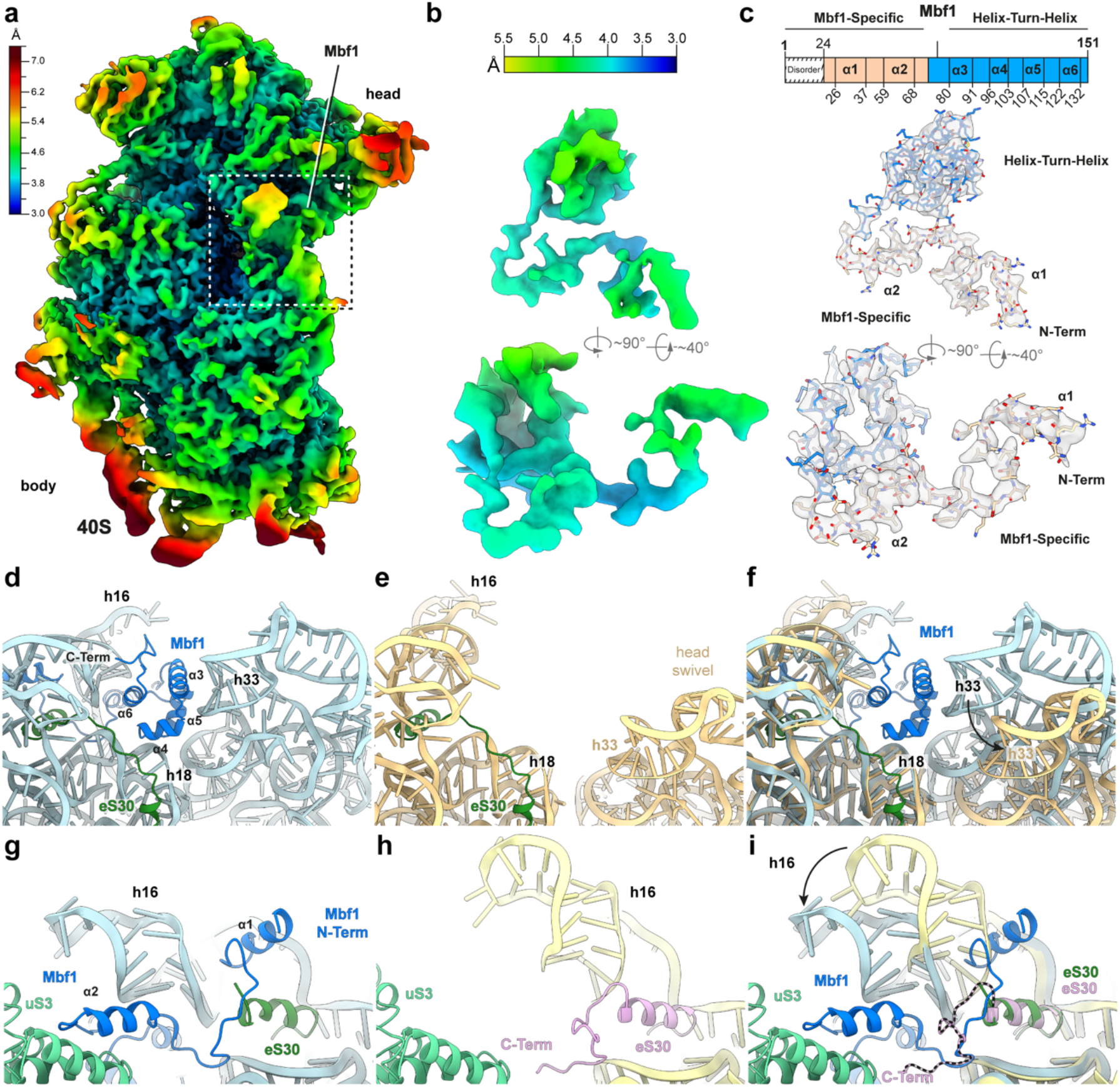
Molecular model for Mbf1 on the Gcn1-disome. (**a**) Local resolution map of the 40S subunit of the colliding ribosome from the Gcn1-disome with the binding site of Mbf1 indicated. (**b**) Two views of the isolated density of the Mbf1 colored according to local resolution. (**c**) Schematic representation of Mbf1 showing the Mbf1-specific N-terminus (light brown, residues 44-79) and the C-terminal helix-turn-helix domain (blue). Isolated density (grey transparency) with fitted molecular model for Mbf1 (colored by domain as illustrated in the scheme). (**d**) Mbf1 C-terminal helix-turn-helix interacting with the 18S rRNA. (**e**) Head swivel 40S ribosome (yellow). (**f**) Similar view to (**d** and **e**) with an overlay of head swivel 40S ribosome (yellow). (**g-i**) Conformation change of eS30 C-terminal region and h16 in the Mbf1 bound structure compared to a reference structure (PDB ID 6SNT, pink for eS30 and yellow for h16).

**Fig. S9.**
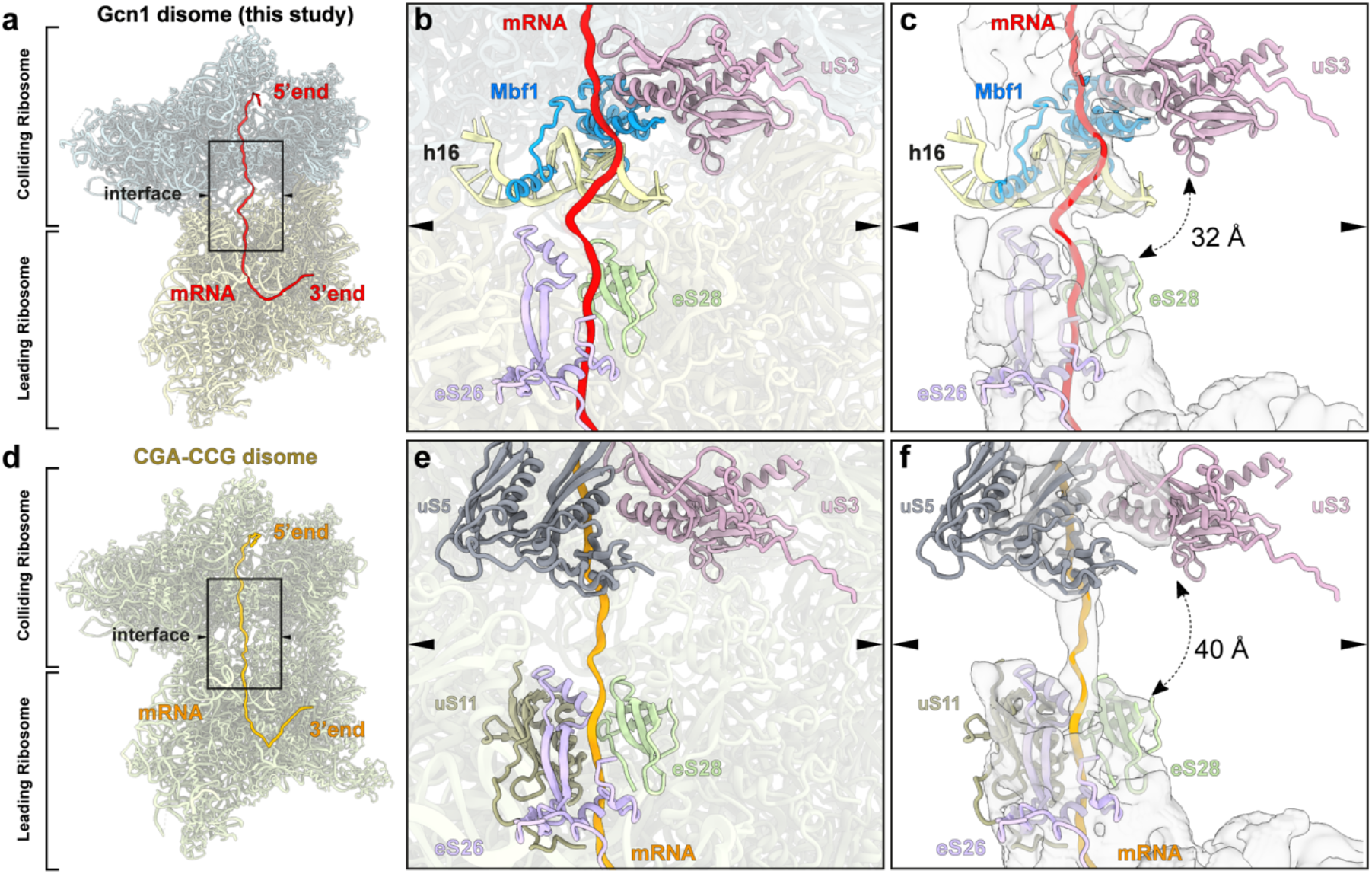
Compaction of the Gcn1-disome. (**a**) 40S of the leading (pale yellow) and the colliding (cyan) ribosome of the Gcn1-stalled disome containing the mRNA (red). (**b**) Close-up view of the 40S-40S interface shown in (**a**). The ribosomal proteins and rRNA interacting with the mRNA are depicted and labelled. (**c**) View as in (**b**) including the locally isolated density for the mRNA and the surrounding ribosomal components. (**d**) 40S subunits (tan) of the disome stalled on CGA-CCG mRNA (PDB ID: 6i7o) (Ikeuchi et al., 2019) (mRNA in light orange) and its (**e**) enlarged view of the 40S-40S interface (**f**) including the locally isolated density for the mRNA and the surrounding ribosomal proteins.

**Table S1.** Cryo-EM data collection, refinement and validation statistics.

|  | Leading stalled ribosome<br>EMD ID XXX<br>PDB ID XXXX | Colliding ribosome<br>EMD ID XXX<br>PDB ID XXXX |
| --- | --- | --- |
| **Data collection** |  |  |
| Microscope | FEI Titan Krios | FEI Titan Krios |
| Camera | Falcon II | Falcon II |
| Magnification | 131,703 | 131,703 |
| Voltage (kV) | 300 | 300 |
| Electron dose (e<sup>-</sup>/Å<sup>2</sup>) | 25 | 25 |
| Defocus range (μm) | -1.3 to -2.8 | -1.3 to -2.8 |
| Pixel size (Å) | 1.084 | 1.084 |
| Initial particles (no.) | 616,079 | 616,079 |
| Final particles (no.) | 30,016 | 30,016 |
| Map resolution (Å) | 3.85 | 4.36 |
| Box size | 700,700,700 | 700,700,700 |
| **Model composition** |  |  |
| Protein residues | 11829 | 11163 |
| RNA bases | 5391 | 5430 |
| **Refinement** |  |  |
| Resolution range (Å) | 3.0-15 | 3.3-15 |
| Map CC (around atoms) | 0.74 | 0.74 |
| Map sharpening *B* factor (Å<sup>2</sup>) | -84.4315 | -120.184 |
| R.m.s. deviations |  |  |
| Bond lengths (Å) (#>4σ) | 0.006 | 0.012 |
| Bond angles (°) (#>4σ) | 1.311 | 1.384 |
| **Validation** |  |  |
| MolProbity score | 1.76 | 2.06 |
| Clashscore | 4.43 | 9.22 |
| Poor rotamers (%) | 0.27 | 1.21 |
| **Ramachandran plot** |  |  |
| Favored (%) | 90.44 | 89.29 |
| Allowed (%) | 9.46 | 10.50 |
| Disallowed (%) | 0.09 | 0.16 |

## References

Adams, P.D., Afonine, P.V., Bunkoczi, G., Chen, V.B., Davis, I.W., Echols, N., Headd, J.J., Hung, L.W., Kapral, G.J., Grosse-Kunstleve, R.W., et al. (2010). PHENIX: a comprehensive Python-based system for macromolecular structure solution. Acta crystallographica. Section D, Biological crystallography 66, 213–221.

Anda, S., Zach, R., and Grallert, B. (2017). Activation of Gcn2 in response to different stresses. PLoS One 12, e0182143.

Andersen, C.B., Becker, T., Blau, M., Anand, M., Halic, M., Balar, B., Mielke, T., Boesen, T., Pedersen, J.S., Spahn, C.M., et al. (2006). Structure of eEF3 and the mechanism of transfer RNA release from the E-site. Nature 443, 663–668.

Andrade, M.A., Petosa, C., O’Donoghue, S.I., Muller, C.W., and Bork, P. (2001). Comparison of ARM and HEAT protein repeats. J Mol Biol 309, 1–18.

Anger, A.M., Armache, J.P., Berninghausen, O., Habeck, M., Subklewe, M., Wilson, D.N., and Beckmann, R. (2013). Structures of the human and Drosophila 80S ribosome. Nature 497, 80–85.

Behrmann, E., Loerke, J., Budkevich, T.V., Yamamoto, K., Schmidt, A., Penczek, P.A., Vos, M.R., Burger, J., Mielke, T., Scheerer, P., et al. (2015). Structural snapshots of actively translating human ribosomes. Cell 161, 845–857.

Bienert, S., Waterhouse, A., de Beer, T.A., Tauriello, G., Studer, G., Bordoli, L., and Schwede, T. (2017). The SWISS-MODEL Repository-new features and functionality. Nucleic acids research 45, D313–D319.

Brandman, O., and Hegde, R.S. (2016). Ribosome-associated protein quality control. Nature structural & molecular biology 23, 7–15.

Brown, A., Baird, M.R., Yip, M.C., Murray, J., and Shao, S. (2018). Structures of translationally inactive mammalian ribosomes. eLife 7.

Castilho, B.A., Shanmugam, R., Silva, R.C., Ramesh, R., Himme, B.M., and Sattlegger, E. (2014). Keeping the eIF2 alpha kinase Gcn2 in check. Biochim Biophys Acta 1843, 1948–1968.

Croll, T.I. (2018). ISOLDE: a physically realistic environment for model building into low-resolution electron-density maps. Acta Crystallogr D Struct Biol 74, 519–530.

Darnell, A.M., Subramaniam, A.R., and O’Shea, E.K. (2018). Translational Control through Differential Ribosome Pausing during Amino Acid Limitation in Mammalian Cells. Molecular cell 71, 229–243 e211.

Daugeron, M.C., Prouteau, M., Lacroute, F., and Seraphin, B. (2011). The highly conserved eukaryotic DRG factors are required for efficient translation in a manner redundant with the putative RNA helicase Slh1. Nucleic acids research 39, 2221–2233.

Deng, J., Harding, H.P., Raught, B., Gingras, A.C., Berlanga, J.J., Scheuner, D., Kaufman, R.J., Ron, D., and Sonenberg, N. (2002). Activation of GCN2 in UV-irradiated cells inhibits translation. Curr Biol 12, 1279–1286.

Elias, J.E., and Gygi, S.P. (2007). Target-decoy search strategy for increased confidence in large-scale protein identifications by mass spectrometry. Nature methods 4, 207–214.

Emsley, P., and Cowtan, K. (2004). Coot: model-building tools for molecular graphics. Acta crystallographica. Section D, Biological crystallography 60, 2126–2132.

Francis, S.M., Gas, M.E., Daugeron, M.C., Bravo, J., and Seraphin, B. (2012). Rbg1-Tma46 dimer structure reveals new functional domains and their role in polysome recruitment. Nucleic acids research 40, 11100–11114.

Goddard, T.D., Huang, C.C., Meng, E.C., Pettersen, E.F., Couch, G.S., Morris, J.H., and Ferrin, T.E. (2018). UCSF ChimeraX: Meeting modern challenges in visualization and analysis. Protein science : a publication of the Protein Society 27, 14–25.

Gutierrez, E., Shin, B.S., Woolstenhulme, C.J., Kim, J.R., Saini, P., Buskirk, A.R., and Dever, T.E. (2013). eIF5A promotes translation of polyproline motifs. Molecular cell 51, 35–45.

Harding, H.P., Ordonez, A., Allen, F., Parts, L., Inglis, A.J., Williams, R.L., and Ron, D. (2019). The ribosomal P-stalk couples amino acid starvation to GCN2 activation in mammalian cells. eLife 8.

Hinnebusch, A.G. (2005). Translational regulation of GCN4 and the general amino acid control of yeast. Annu Rev Microbiol 59, 407–450.

Ikeuchi, K., Tesina, P., Matsuo, Y., Sugiyama, T., Cheng, J., Saeki, Y., Tanaka, K., Becker, T., Beckmann, R., and Inada, T. (2019). Collided ribosomes form a unique structural interface to induce Hel2-driven quality control pathways. EMBO J 38.

Inada, T. (2020). Quality controls induced by aberrant translation. Nucleic acids research 48, 1084–1096.

Ishikawa, K., Akiyama, T., Ito, K., Semba, K., and Inoue, J. (2009). Independent stabilizations of polysomal Drg1/Dfrp1 complex and non-polysomal Drg2/Dfrp2 complex in mammalian cells. Biochem Biophys Res Commun 390, 552–556.

Ishikawa, K., Azuma, S., Ikawa, S., Semba, K., and Inoue, J. (2005). Identification of DRG family regulatory proteins (DFRPs): specific regulation of DRG1 and DRG2. Genes Cells 10, 139–150.

Ishikawa, K., Ito, K., Inoue, J., and Semba, K. (2013). Cell growth control by stable Rbg2/Gir2 complex formation under amino acid starvation. Genes Cells 18, 859–872.

Ishimura, R., Nagy, G., Dotu, I., Chuang, J.H., and Ackerman, S.L. (2016). Activation of GCN2 kinase by ribosome stalling links translation elongation with translation initiation. eLife 5.

Joazeiro, C.A.P. (2019). Mechanisms and functions of ribosome-associated protein quality control. Nat Rev Mol Cell Biol 20, 368–383.

Juszkiewicz, S., Chandrasekaran, V., Lin, Z., Kraatz, S., Ramakrishnan, V., and Hegde, R.S. (2018). ZNF598 Is a Quality Control Sensor of Collided Ribosomes. Molecular cell 72, 469–481 e467.

Juszkiewicz, S., Slodkowicz, G., Lin, Z., Freire-Pritchett, P., Peak-Chew, S.Y., and Hegde, R.S. (2020a). Ribosome collisions trigger cis-acting feedback inhibition of translation initiation. eLife 9.

Juszkiewicz, S., Speldewinde, S.H., Wan, L., Svejstrup, J.Q., and Hegde, R.S. (2020b). The ASC-1 Complex Disassembles Collided Ribosomes. Molecular cell 79, 603–614 e608.

Kubota, H., Ota, K., Sakaki, Y., and Ito, T. (2001). Budding yeast GCN1 binds the GI domain to activate the eIF2alpha kinase GCN2. The Journal of biological chemistry 276, 17591–17596.

Kubota, H., Sakaki, Y., and Ito, T. (2000). GI domain-mediated association of the eukaryotic initiation factor 2alpha kinase GCN2 with its activator GCN1 is required for general amino acid control in budding yeast. The Journal of biological chemistry 275, 20243–20246.

Lee, S.J., Swanson, M.J., and Sattlegger, E. (2015). Gcn1 contacts the small ribosomal protein Rps10, which is required for full activation of the protein kinase Gcn2. Biochem J 466, 547–559.

Marton, M.J., Crouch, D., and Hinnebusch, A.G. (1993). GCN1, a translational activator of GCN4 in Saccharomyces cerevisiae, is required for phosphorylation of eukaryotic translation initiation factor 2 by protein kinase GCN2. Molecular and cellular biology 13, 3541–3556.

Marton, M.J., Vazquez de Aldana, C.R., Qiu, H., Chakraburtty, K., and Hinnebusch, A.G. (1997). Evidence that GCN1 and GCN20, translational regulators of GCN4, function on elongating ribosomes in activation of eIF2alpha kinase GCN2. Molecular and cellular biology 17, 4474–4489.

Matsuo, Y., Tesina, P., Nakajima, S., Mizuno, M., Endo, A., Buschauer, R., Cheng, J., Shounai, O., Ikeuchi, K., Saeki, Y., et al. (2020). RQT complex dissociates ribosomes collided on endogenous RQC substrate SDD1. Nature structural & molecular biology 27, 323–332.

Meydan, S., and Guydosh, N.R. (2020). Disome and Trisome Profiling Reveal Genome-wide Targets of Ribosome Quality Control. Molecular cell 79, 588–602 e586.

Pettersen, E.F., Goddard, T.D., Huang, C.C., Couch, G.S., Greenblatt, D.M., Meng, E.C., and Ferrin, T.E. (2004). UCSF Chimera--a visualization system for exploratory research and analysis. Journal of computational chemistry 25, 1605–1612.

Pittman, Y.R., Valente, L., Jeppesen, M.G., Andersen, G.R., Patel, S., and Kinzy, T.G. (2006). Mg2+ and a key lysine modulate exchange activity of eukaryotic translation elongation factor 1B alpha. The Journal of biological chemistry 281, 19457–19468.

Salinas, R.K., Camilo, C.M., Tomaselli, S., Valencia, E.Y., Farah, C.S., El-Dorry, H., and Chambergo, F.S. (2009). Solution structure of the C-terminal domain of multiprotein bridging factor 1 (MBF1) of Trichoderma reesei. Proteins 75, 518–523.

Sattlegger, E., and Hinnebusch, A.G. (2000). Separate domains in GCN1 for binding protein kinase GCN2 and ribosomes are required for GCN2 activation in amino acid-starved cells. EMBO J 19, 6622–6633.

Sattlegger, E., and Hinnebusch, A.G. (2005). Polyribosome binding by GCN1 is required for full activation of eukaryotic translation initiation factor 2{alpha} kinase GCN2 during amino acid starvation. The Journal of biological chemistry 280, 16514–16521.

Schmidt, C., Becker, T., Heuer, A., Braunger, K., Shanmuganathan, V., Pech, M., Berninghausen, O., Wilson, D.N., and Beckmann, R. (2016a). Structure of the hypusinylated eukaryotic translation factor eIF-5A bound to the ribosome. Nucleic acids research 44, 1944–1951.

Schmidt, C., Kowalinski, E., Shanmuganathan, V., Defenouillere, Q., Braunger, K., Heuer, A., Pech, M., Namane, A., Berninghausen, O., Fromont-Racine, M., et al. (2016b). The cryo-EM structure of a ribosome-Ski2-Ski3-Ski8 helicase complex. Science 354, 1431–1433.

Schuller, A.P., Wu, C.C., Dever, T.E., Buskirk, A.R., and Green, R. (2017). eIF5A Functions Globally in Translation Elongation and Termination. Molecular cell 66, 194–205 e195.

Schwanhausser, B., Busse, D., Li, N., Dittmar, G., Schuchhardt, J., Wolf, J., Chen, W., and Selbach, M. (2011). Global quantification of mammalian gene expression control. Nature 473, 337–342.

Simms, C.L., Yan, L.L., Qiu, J.K., and Zaher, H.S. (2019). Ribosome Collisions Result in +1 Frameshifting in the Absence of No-Go Decay. Cell reports 28, 1679–1689 e1674.

Sinha, N.K., Ordureau, A., Best, K., Saba, J.A., Zinshteyn, B., Sundaramoorthy, E., Fulzele, A., Garshott, D.M., Denk, T., Thoms, M., et al. (2020). EDF1 coordinates cellular responses to ribosome collisions. eLife 9.

Svidritskiy, E., Brilot, A.F., Koh, C.S., Grigorieff, N., and Korostelev, A.A. (2014). Structures of yeast 80S ribosome-tRNA complexes in the rotated and nonrotated conformations. Structure 22, 1210–1218.

Tesina, P., Lessen, L.N., Buschauer, R., Cheng, J., Wu, C.C., Berninghausen, O., Buskirk, A.R., Becker, T., Beckmann, R., and Green, R. (2020). Molecular mechanism of translational stalling by inhibitory codon combinations and poly(A) tracts. EMBO J 39, e103365.

Triana-Alonso, F.J., Chakraburtty, K., and Nierhaus, K.H. (1995). The elongation factor 3 unique in higher fungi and essential for protein biosynthesis is an E site factor. The Journal of biological chemistry 270, 20473–20478.

Tyanova, S., Temu, T., and Cox, J. (2016). The MaxQuant computational platform for mass spectrometry-based shotgun proteomics. Nat Protoc 11, 2301–2319.

Vazquez de Aldana, C.R., Marton, M.J., and Hinnebusch, A.G. (1995). GCN20, a novel ATP binding cassette protein, and GCN1 reside in a complex that mediates activation of the eIF-2 alpha kinase GCN2 in amino acid-starved cells. EMBO J 14, 3184–3199.

Visweswaraiah, J., Lee, S.J., Hinnebusch, A.G., and Sattlegger, E. (2012). Overexpression of eukaryotic translation elongation factor 3 impairs Gcn2 protein activation. The Journal of biological chemistry 287, 37757–37768.

Wang, J., Zhou, J., Yang, Q., and Grayhack, E.J. (2018). Multi-protein bridging factor 1(Mbf1), Rps3 and Asc1 prevent stalled ribosomes from frameshifting. eLife 7.

Wolf, A.S., and Grayhack, E.J. (2015). Asc1, homolog of human RACK1, prevents frameshifting in yeast by ribosomes stalled at CGA codon repeats. RNA 21, 935–945.

Wout, P.K., Sattlegger, E., Sullivan, S.M., and Maddock, J.R. (2009). Saccharomyces cerevisiae Rbg1 protein and its binding partner Gir2 interact on Polyribosomes with Gcn1. Eukaryot Cell 8, 1061–1071.

Wu, C.C., Peterson, A., Zinshteyn, B., Regot, S., and Green, R. (2020). Ribosome Collisions Trigger General Stress Responses to Regulate Cell Fate. Cell 182, 404–416 e414.

Zeng, F., Pires-Alves, M., Hawk, C.W., Chen, X., and Jin, H. (2020). Molecular Functions of Conserved Developmentally-Regulated GTP-Binding Protein Drg1 in Translation. BioRxiv.

Zivanov, J., Nakane, T., Forsberg, B.O., Kimanius, D., Hagen, W.J., Lindahl, E., and Scheres, S.H. (2018). New tools for automated high-resolution cryo-EM structure determination in RELION-3. eLife 7.

